# Mitochondrial collapse links PFKFB3-promoted glycolysis with CLN7/MFSD8 neuronal ceroid lipofuscinosis pathogenesis

**DOI:** 10.1101/2020.10.20.345314

**Authors:** Irene Lopez-Fabuel, Marina Garcia-Macia, Costantina Buondelmonte, Olga Burmistrova, Nicolo Bonora, Brenda Morant-Ferrando, Paula Alonso-Batan, Carlos Vicente-Gutierrez, Daniel Jimenez-Blasco, Ruben Quintana-Cabrera, Emilio Fernandez, Aseel Sharaireh, Marta Guevara-Ferrer, Lorna Fitzpatrick, Christopher D. Thompton, Tristan R. McKay, Stephan Storch, Diego L. Medina, Sara E. Mole, Peter O. Fedichev, Angeles Almeida, Juan P. Bolaños

**Affiliations:** Institute of Functional Biology and Genomics (IBFG), Universidad de Salamanca, CSIC, Salamanca, Spain; Institute of Biomedical Research of Salamanca (IBSAL), Hospital Universitario de Salamanca, CSIC, Universidad de Salamanca, Salamanca, Spain; Centro de Investigación Biomédica en Red de Fragilidad y Envejecimiento Saludable (CIBERFES), Madrid, Spain; Gero Discovery LLC, Moscow, Russia; Centre for Bioscience, Manchester Metropolitan University, Manchester M1 5GD, United Kingdom; University Children’s Research@Kinder-UKE, University Medical Center Hamburg-Eppendorf, Hamburg, Germany; Telethon Institute of Genetics and Medicine (TIGEM), High Content Screening Facility, Via Campi Flegrei 34, 80078 Pozzuoli, Italy; and Medical Genetics Unit, Department of Medical and Translational Science, Federico II University, 80138 Naples, Italy; MRC Laboratory for Molecular Biology and GOS Institute for Child Health, University College London, United Kingdom

## Abstract

The neuronal ceroid lipofuscinoses (NCLs) are a family of monogenic life-limiting pediatric neurodegenerative disorders collectively known as Batten disease^1^. Although genetically heterogeneous^2^, NCLs share several clinical symptoms and pathological hallmarks such as lysosomal accumulation of lipofuscin and astrogliosis^2,3^. CLN7 disease belongs to a group of NCLs that present in late infancy^4–6^ and, whereas *CLN7/MFSD8* gene is known to encode a lysosomal membrane glycoprotein^4,7–9^, the biochemical processes affected by CLN7-loss of function are unexplored thus preventing development of potential treatments^1,10^. Here, we found in the *Cln7*^Δ*ex2*^ mouse model^11^ of CLN7 disease that failure in the autophagy-lysosomal pathway causes accumulation of structurally and bioenergetically impaired, reactive oxygen species (ROS)-producing neuronal mitochondria that contribute to CLN7 pathogenesis. *Cln7*^Δ*ex2*^ neurons exhibit a metabolic shift mediated by pro-glycolytic enzyme 6-phosphofructo-2-kinase/fructose-2,6-bisphosphatase-3 (PFKFB3). PFKFB3 inhibition in *Cln7*^Δ*ex2*^ mice *in vivo* and in CLN7 patients-derived cells rectified key disease hallmarks. Thus, specifically targeting glycolysis may alleviate CLN7 pathogenesis.

---

Forty-six disease-causing mutations are recorded in the NCL mutation database (ucl.ac.uk/ncl-disease) in *CLN7/MFSD8*, causing a broad phenotypic range, from classic late infantile CLN7 disease to non-syndromic retinal disease with onset in childhood or as late as the 7^th^ decade^4^. Given that treatment for CLN7 disease is likely to be more challenging than for NCLs encoding lysosomal enzymes such as CLN2/TPP1 (Ref. 12), here we aimed to understand the biochemical processes affected in CLN7 disease. In Cln7-null neurons in primary culture from *Cln7*^Δ*ex2*^ mice^11^ (**Extended Data Fig. 1a**), the mitochondrial indicators ATP synthase-subunit c (SCMAS) and heat-shock protein-60 (HSP60) co-localized with the lysosome-associated membrane protein-1 (LAMP1) (**Fig. 1a; Extended Data Fig. 1b**), suggesting lysosomal accumulation of mitochondria. To determine mitochondrial turnover, neurons were incubated with inhibitors of lysosomal proteolysis, a treatment that triggered the accumulation of SCMAS and HSP60 in wild type (WT) neurons (**Fig 1b; Extended Data Fig. 1c**), indicating mitophagy flux^13^. However, these mitochondrial markers were already increased in untreated *Cln7*^Δ*ex2*^ neurons and were little affected by inhibiting lysosomal function (**Fig. 1b; Extended Data Fig. 1c**). In addition, PTEN-induced kinase-1 (PINK1) 63/53 ratio^14^ increased in *Cln7*^Δ*ex2*^ neurons (**Extended Data Fig. 1d**). These data suggest that the mitochondrial clearance in *Cln7*^Δ*ex2*^ neurons is impaired. The metabolic profile analysis revealed a decrease in the basal oxygen consumption rate (OCR), ATP-linked and maximal OCR, and proton leak in *Cln7*^Δ*ex2*^ neurons (**Fig. 1c**), indicating bioenergetically impaired mitochondria. The specific activities of the mitochondrial respiratory chain (MRC) complexes (**Extended Data Fig. 1e**) were unchanged in the *Cln7*^Δ*ex2*^ neurons. However, isolation of mitochondria followed by blue-native gel electrophoresis (BNGE), complex I (CI) in-gel activity assay (IGA) and western blotting, revealed CI disassembly from mitochondrial supercomplexes (SCs) in *Cln7*^Δ*ex2*^ neurons (**Fig. 1d**). These data confirm the decreased mitochondrial energy efficiency^15^ and suggest increased formation of mitochondrial reactive oxygen species (mROS)^16^ in *Cln7*^Δ*ex2*^ neurons. Flow cytometric analysis of mROS (**Fig. 1e; Extended Data Fig. 1f**) and fluorescence analysis of H_2_O_2_ (**Extended Data Fig. 1g**) confirmed mROS enhancement in *Cln7*^Δ*ex2*^ neurons. To characterize mitochondria from *Cln7*^Δ*ex2*^ mice *in vivo*, we next performed electron microscopy analyses of the brain cortex before and after the onset of the immunohistochemical and behavioral symptoms of the disease^11^. We found larger and longer brain mitochondria in the pre-symptomatic *Cln7*^Δ*ex2*^ mice, an effect that proceeded with age (**Fig. 1f**; **Extended Data Fig. 1h**), suggesting progressive mitochondrial swelling. CI disassembly from SCs in brain mitochondria (**Fig. 1g**) and increased mROS in freshly purified neurons (**Fig. 1h; Extended Data Fig. 1i**) were confirmed in *Cln7*^Δ*ex2*^ mice. Altogether, these findings suggest that Cln7 loss causes impaired autophagic clearance of brain mitochondria leading to the aberrant accumulation of structurally disorganized, bioenergetically impaired and high ROS-generating organelle.

**Fig. 1.**
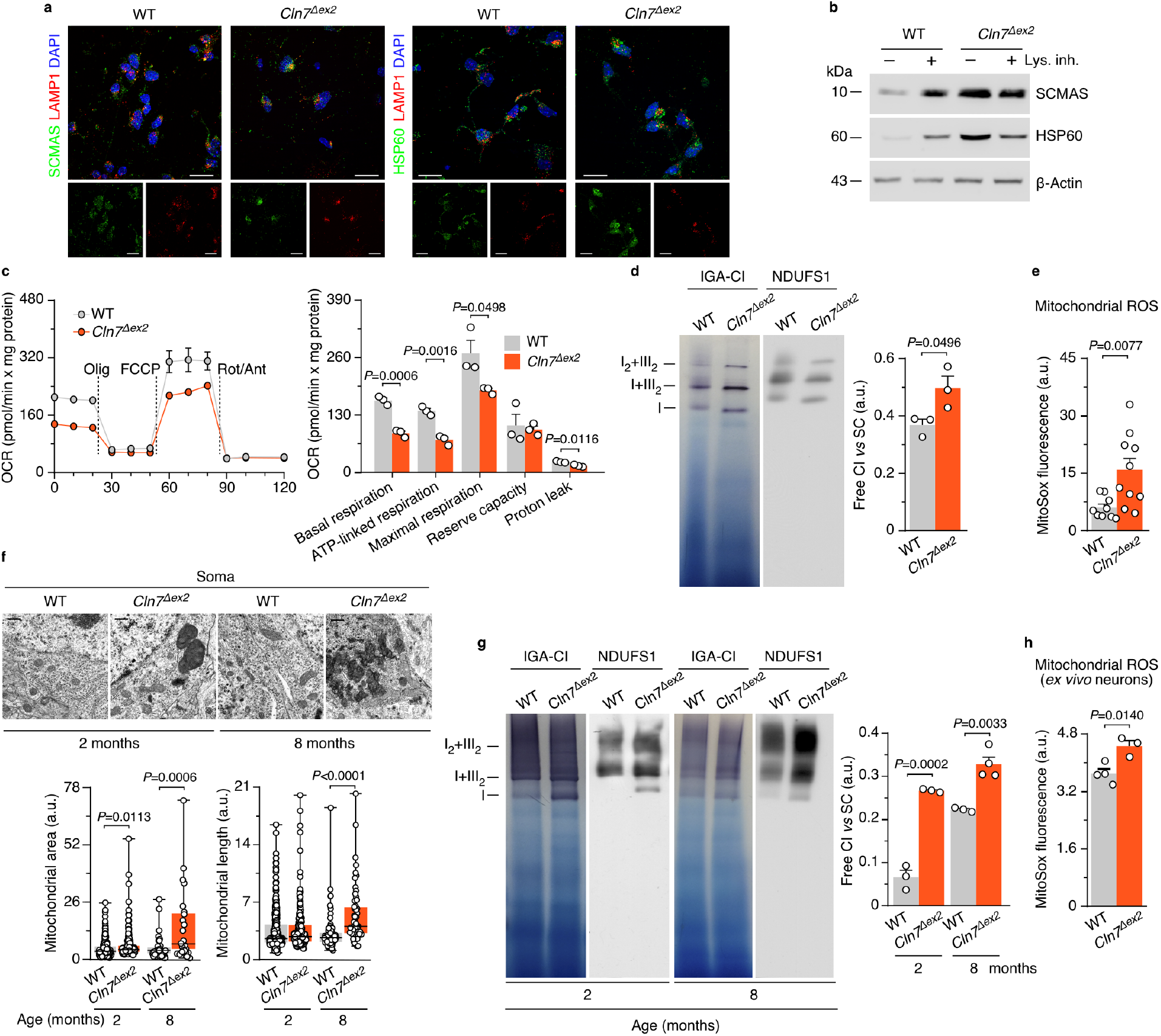
Failure in the autophagy-lysosomal pathway causes accumulation of structural and functionally impaired mitochondria in *Cln7*^Δ*ex2*^ mouse. **(a)** SCMAS/LAMP1 and HSP60/SCMAS co-localization confocal analyses in primary neurons. DAPI reveals nuclei. Scale bar, 20 μm. **(b)** HSP60 and SCMAS western blot analysis in primary neurons incubated with lysosomal proteolysis inhibitors leupeptin (100 μM) plus NH4Cl (20 mM) (Lys. Inh.) for 1 h (ß-actin, loading control). **(c)** OCR analysis (left) and calculated parameters (right) in primary neurons (n=3). **(d)** Free complex I (CI) and CI-containing supercomplexes (SC) analyses in primary neurons by BNGE in-gel activity (IGA-CI) and by immunoblotted PVDF membranes against CI subunit, NDUFS1 (n=3). **(e)** Mitochondrial ROS analysis in primary neurons (n=9-10). **(f)** Representative electron microscopy images and analyses of mouse brain cortex mitochondria (~175 mitochondria per condition). Scale bar, 500 nm. **(g)** Free CI and CI-containing SC analyses of mouse brain cortex by BNGE IGA-CI and by immunoblotted PVDF membranes against NDUFS1 (n=3-4). **(h)** Mitochondrial ROS analysis in freshly isolated mouse brain cortex neurons (n=3-4 of 6-months old mice). Data are mean ± S.E.M. for the indicated number of culture preparations or mice. Statistical analyses were performed by Student’s *t* test. Representative images and western blots out of n≥3 experiments are shown (see Extended Data Fig. 1).

Next, we assessed the impact of excess neuronal mROS on CLN7 disease progression. *Cln7*^Δ*ex2*^ mice were thus crossed with mice expressing a mitochondrial tagged isoform of the H_2_O_2_-detoxifying enzyme catalase (mCAT) governed by the neuron-specific^17^ calcium/calmodulin-dependent protein kinase II alpha (CaMKIIa) promoter (*CaMKIIa^Cre^-mCAT^LoxP^*). mCAT efficacy *in vivo* was previously validated^18^. The resulting progeny (*Cln7^Δex2^-CAMKIIa^Cre^-mCAT*) was analyzed and compared with littermate *Cln7^Δex2^-mCAT^LoxP^* and control (*mCAT^Lop^* and *CAMKIIa^Cre^-mCAT)* mice. The increased mROS observed in *Cln7^Δex2^-mCAT^LoxP^* neurons was abolished in *Cln7^Δex2^-CAMKIIa^Cre^-mCAT* neurons (**Fig. 2a**; **Extended Data Fig. 2a**), verifying the efficacy of this approach. Brain mitochondrial swelling was confirmed in *Cln7^Δex2^-mCAT^LoxP^* mice (**Fig. 2b**), which also showed mitochondrial cristae profile widening (**Fig. 2b**), a phenomenon previously observed in cells with bioenergetically-inefficient mitochondria^19,20^. Both the mitochondrial swelling and cristae profile widening observed in the (*Cln7Γ^Δ^-mCAT^LoP^* mice were rescued by expressing mCAT in neurons of the *Cln7^Δex2^-CAMKIIa^Cre^-mCAT* mice (**Fig. 2b**). Thus, neuronal mROS participates in the accumulation of functional and ultrastructural impaired mitochondria in *Cln7*^Δ*ex2*^ mouse brain. In line with this, the increase in SCMAS abundance observed in the *Cln7^Δex2^-mCA T^LoxP^* mouse brain (**Fig. 2c; Extended Data Fig. 2b**) was abolished, or partially restored, in *Cln7^Δex2^-CAMKIIa^Cre^-mCAT* mice (**Fig. 2c**; **Extended Data Fig. 2b**). Brain SCMAS accumulation in autofluorescent ceroid lipopigments (lipofuscin)-containing lysosomes is a hallmark of Batten disease^2^. Consistently with this notion, lipofuscin was accumulated in the brain of *Cln7^Δex2^-mCAT^LoxP^* mice, an effect that was ameliorated in *Cln7^Δex2^-CAMKIIa^Cre^-mCAT* mice (**Fig. 2c**; **Extended Data Fig. 2c**). Moreover, activation of astrocytes and microglia is another hallmark of Batten disease^3^ that is mimicked in the brain of *Cln7*^Δ*ex2*^ mice^11^. We found increased glial-fibrillary acidic protein (GFAP) and ionized calcium binding adaptor molecule-1 (IBA-1) proteins in the brain of *Cln7^Δex2^-mCAT^LoxP^* mice, suggesting astrocytosis and microgliosis, respectively; these effects were attenuated in *Cln7^Δex2^-CAMKIIa^Cre^-mCAT* mice (**Fig. 2c**; **Extended Data Fig. 2d**). Altogether, these findings indicate that the generation of ROS by bioenergetically-impaired mitochondria in *Cln7*^Δ*ex2*^ neurons contributes to the histopathological symptoms of CLN7 disease.

**Fig. 2.**
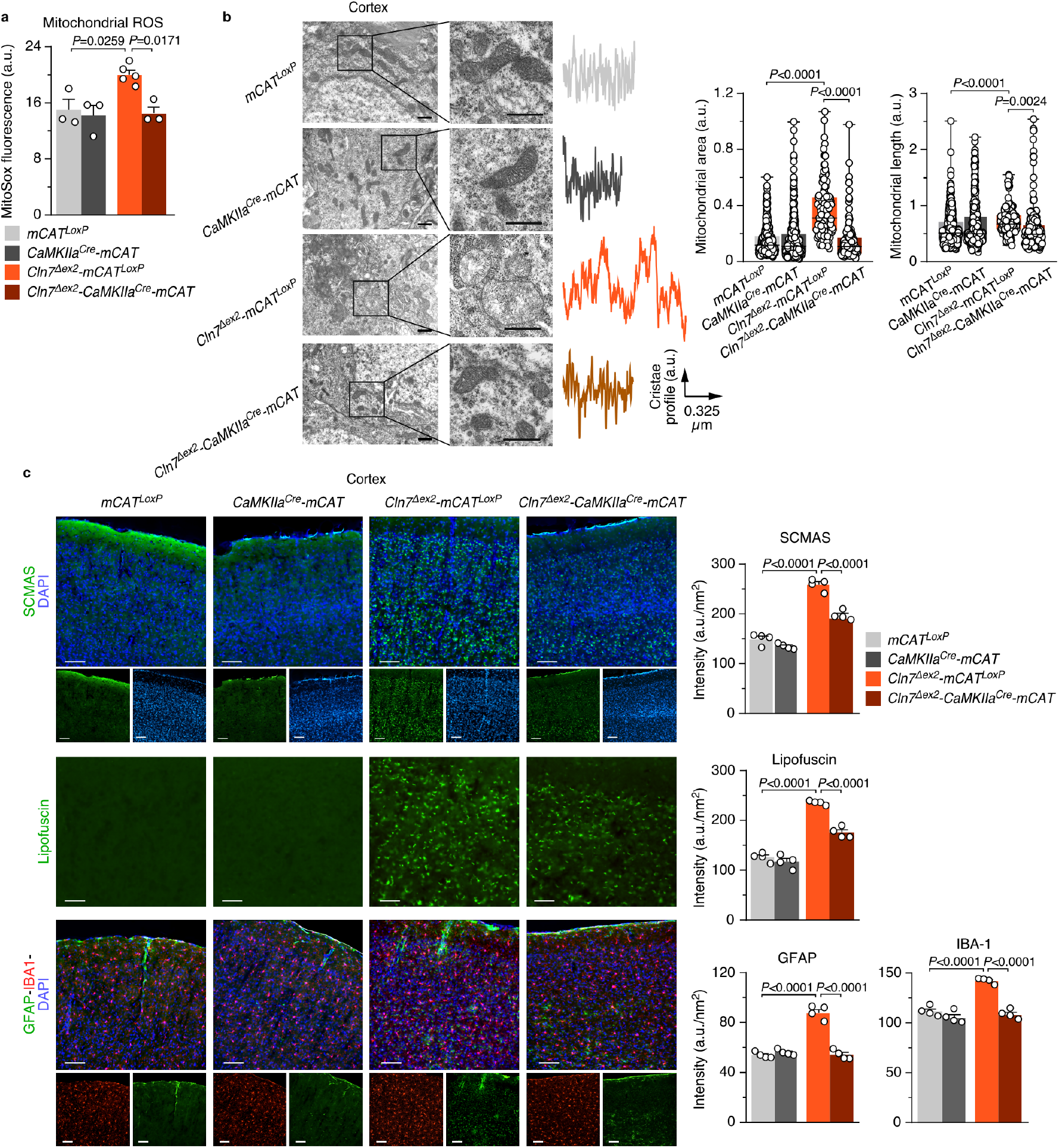
Increased generation of mitochondrial ROS by neurons accounts for impaired mitochondrial accumulation and hallmarks of CLN7 disease in *Cln7*^Δ*ex2*^ mouse *in vivo.* **(a)** Mitochondrial ROS analysis in primary neurons from the designed genotype (n=3-5). **(b)** Representative electron microscopy images and analyses of the mouse brain cortex displaying the cristae profile plot of intensities over the maximal axis of the magnified shown mitochondrion (~290 mitochondria per condition; 3-months old mice). Scale bars, 600 nm. **(c)** SCMAS, lipofuscin, GFAP and IBA-1 immunohistochemical analysis of the mouse brain cortex (3 serial slices per mouse; n=4 of 3-months-old mice). Scale bar, 100 μm. Data are mean ± S.E.M. for the indicated number of culture preparations or mice. Statistical analyses were performed by one-way ANOVA followed by Tukey’s post-hoc test. (See Extended Data Fig. 2).

Mitochondrial ROS stimulate brain glucose consumption through the glycolytic pathway in mouse^18^. In *Cln7^Δex2^-mCAT^LoxP^* neurons, both glycolysis (**Fig. 3a**), and its end-product lactate (**Extended Data Fig. 3a**) were up-regulated, effects that were abolished in *Cln7^Δex2^-CAMKIIa^Cre^-mCAT* neurons. Glycolytic and pentose-phosphate pathway (PPP) fluxes are inversely regulated in neurons^21–23^. Agreeingly, the increased glycolytic flux (**Fig. 3b**) and lactate production (**Extended Data Fig. 3b**) observed in *Cln7*^Δ*ex2*^ neurons was accompanied by reduced PPP flux (**Fig. 3c**). A shift of glucose metabolism from PPP to glycolysis can be indicative of hyperactive 6-phosphofructo-1-kinase (PFK1)^24,25^, a rate-limiting step of glycolysis that is regulated by fructose-2,6-bisphosphate (F-2,6-P2), a robust positive effector of PFK1 (Ref. 26). The rate of F-2,6-P2 formation was enhanced in *Cln7*^Δ*ex2*^ neurons (**Fig. 3d**), a result that is compatible with higher activity of 6-phosphofructo-2-kinase/fructose-2,6-bisphosphatase-3 (PFKFB3) -i.e. the only F-2,6-P2-forming isoenzyme found in neurons upon stress conditions^21^. PFKFB3 protein was increased both in primary neurons and *in vivo* brain cortex and cerebellum (**Fig. 3e; Extended Data Fig. 3c,d**) of the *Cln7*^Δ*ex2*^ mice. Since PFKFB3 mRNA abundance was unaltered in *Cln7*^Δ*ex2*^ neurons (**Fig. 3f**), we conjectured that increased PFKFB3 protein could be consequence of inactivating its degrading pathway^21^. *Cln7*^Δ*ex2*^ neurons showed hyperphosphorylation of the anaphase-promoting complex/cyclosome (APC/C) activator protein, Cdh1 (**Fig. 3g; Extended Data Fig. 3e**), which is sufficient to inhibit APC/C E3-ligase activity that targets PFKFB3 for proteasomal degradation^21^. To pursue this possibility, we noted that the Ca^2+^-buffering capacity of bioenergetically-compromised mitochondria is impaired^27^. Indeed, *Cln7*^Δ*ex2*^ neurons showed enhanced concentration of cytosolic Ca^2−^ (**Fig. 3h**), an activator of calpain -a proteolytic enzyme essential in the signaling cascade leading to Cdh1 hyperphosphorylation^28^. Ca^2+^ sequestration reduced both PFKFB3 protein (**Fig. 3i; Extended Data Fig. 3f**) and glycolysis (**Fig. 3j**) in *Cln7*^Δ*ex2*^ neurons, confirming Ca^2+^ involvement in increasing glycolytic flux. Ca^2+^-mediated calpain activation proteolytically cleaves p35 into p25 -a cofactor of the cyclin-dependent kinase-5 (Cdk5)^29^ that phosphorylates Cdh1 (Ref. 28). We found an increased p35 cleavage into p25 in *Cln7*^Δ*ex2*^ primary neurons and *in vivo* brain cortex and cerebellum (**Fig. 3k; Extended Data Fig. 3g,h**). Inhibition of calpain using the specific inhibitor^29^ MDL-28170 rescued p35 cleavage and PFKFB3 increase (**Fig. 3l; Extended Data Fig. 3i**). Given that these effects suggest the involvement of Cdk5, *Cdk5* was knocked down in *Cln7*^Δ*ex2*^ neurons, an action that prevented PFKFB3 increase (**Fig. 3m; Extended Data Fig. 3j**). Together, these results indicate the occurrence of a Ca^2+^/calpain-mediated activation of Cdk5/p25 pathway that phosphorylates APC/C cofactor Cdh1, eventually leading to the stabilization of pro-glycolytic PFKFB3 in CLN7 disease.

**Fig. 3.**
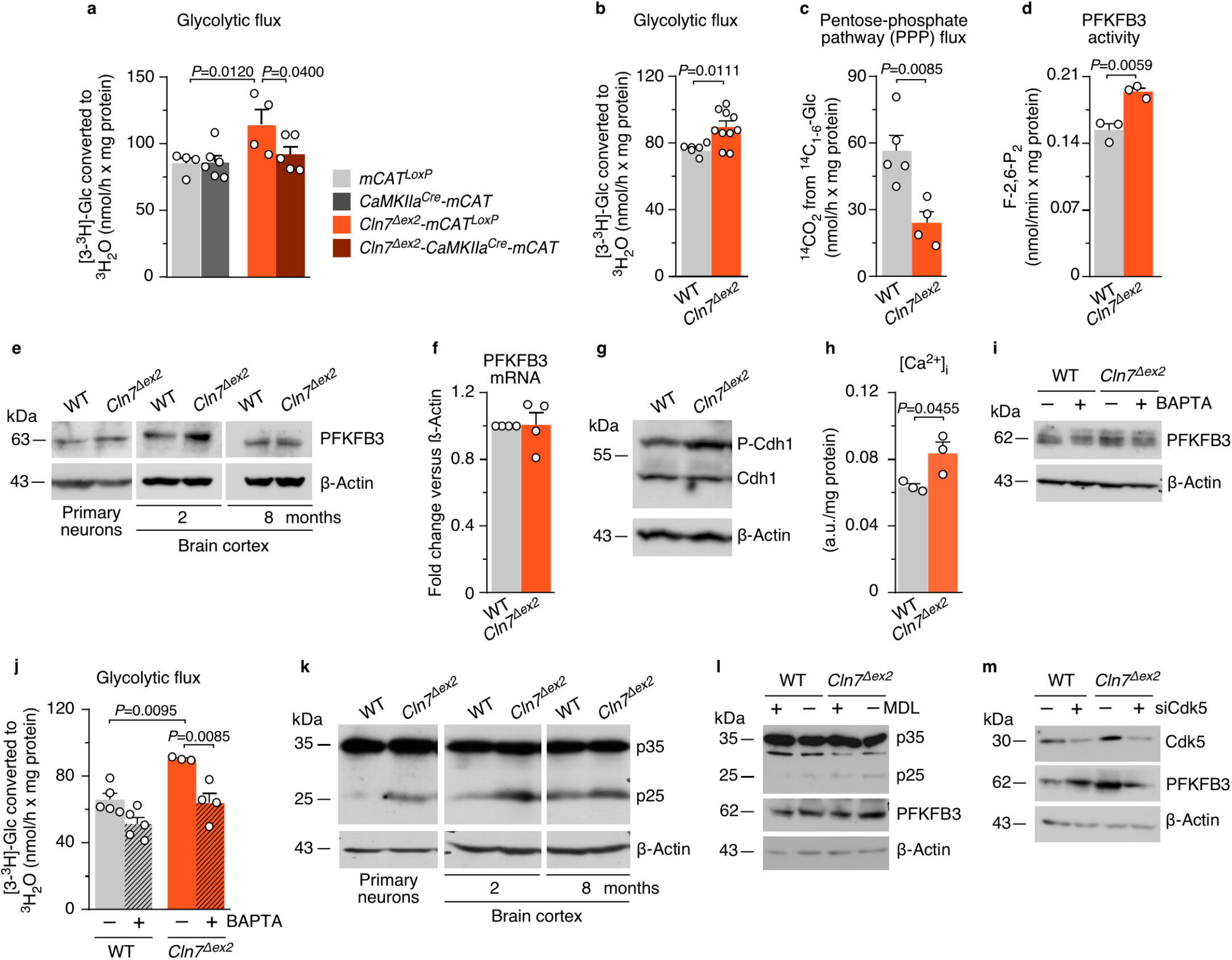
Upregulation of PFKFB3 protein and activity *via* a Ca^2+^/calpain/Cdk5 pathway sustains a high glycolytic flux in *Cln7*^Δ*ex2*^ neurons. **(a,b)** Glycolytic flux in primary neurons (**a**, n=4-5; **b**, n=6-10). **(c)** PPP flux in primary neurons (n=4-5). **(d)** Rate of P-2,6-P2 formation in primary neurons (n=3). **(e)** PFKFB3 western blot analysis in primary neurons and brain cortex (ß-actin, loading control). **(f)** PFKFB3 mRNA analysis by RT-qPCR in primary neurons (n=4; values normalized *versus* ß-actin). **(g)** Cdh1 western blot analysis after PhosTag acrylamide electrophoresis in primary neurons and brain cortex (P-Cdh1, hyperphosphorylated Cdh1; ß-actin, loading control). **(h)** Cytosolic Ca^2+^ analysis in primary neurons (n=3). **(i,j)** PFKFB3 western blot (**i**) and glycolytic flux (**j**) analyses in primary neurons incubated with Ca^2+^ quelator BAPTA (10 μM; 1 h) (ß-actin, loading control) (**j**, n=3-5). **(k)** p35 western blot revealing p35 and its cleavage product p25 in primary neurons and brain cortex (ß-actin, loading control). **(l)** p35 and PFKFB3 western blot analyses in primary neurons incubated with calpain inhibitor MDL-28170 (MDL) (10 μM; 24 h) (ß-actin, loading control). **(m)** Cdk5 and PFKFB3 western blot analyses in primary neurons transfected with Cdk5 siRNA (siCdk5) or scrambled siRNA (−) (9 nM; 3 days) (ß-actin, loading control). Data are mean ± S.E.M. for the indicated number of culture preparations or mice. Statistical analyses performed by one-way ANOVA followed by DMS’s (**a**) or Tukey’s (**j**) post-hoc tests or Student’s t test (**b, c, d, h**). Representative western blots out of n≥3 experiments are shown (see Extended Data Fig. 3).

In neurons, PFKFB3 destabilization boosts glucose consumption through PPP^21^ and prevents damage-associated redox stress^21,30,31^ given its role at supplying NADPH(H^+^) – an essential cofactor of glutathione regeneration^32,33^. We therefore sought to assess whether PFKFB3-promoted glycolysis is related to CLN7 disease. We undertook this by inhibiting PFKFB3 activity using the highly selective, rationally designed^34^ compound AZ67. Incubation of *Cln7*^Δ*ex2*^ neurons with AZ67 at a concentration that inhibits PFKFB3 activity without compromising survival^35^, prevented the increase in F-2,6-P2 (**Fig. 4a**) and glycolysis (**Fig. 4b**) without affecting mROS (**Fig. 4c; Extended Data Fig. 4a**). Interestingly, AZ67 protected *Cln7*^Δ*ex2*^ neurons from activation of pro-apoptotic caspase-3 (**Fig. 4d; Extended Data Fig. 4b**), suggesting its potential therapeutic benefit. To test this *in vivo*, AZ67 was intracerebroventricularly administered in *Cln7*^Δ*ex2*^ mice daily for two months at a previously chosen dose (**Extended Data Fig. 4c,d,e**). Electron microscopy analysis revealed that AZ67 did not affect the length or area of brain mitochondria in *Cln7*^Δ*ex2*^ mice (**Fig. 4e**), but it prevented the cristae profile amplitude reduction (**Fig. 4e**); this may indicate, as observed in other paradigms^19,20^, adaptation of the mitochondrial ultrastructure to a bioenergetically efficient configuration upon glycolysis inhibition. Incubation of *Cln7*^Δ*ex2*^ neurons with AZ67 partially restored the impairment in basal respiration (**Fig. 4f**), indicating a functional improvement of mitochondria. *In vivo*, AZ67 prevented the accumulation of SCMAS, lipofuscin and reactive astroglia in the cortex (**Fig. 4g**), and SCMAS and lipofuscin in the hippocampus and cerebellum (**Extended Data Fig. 5a,b,c**) of the *Cln7*^Δ*ex2*^ mice. Hindlimb paralysis^11^ in *Cln7*^Δ*ex2*^ mice was prevented by AZ67 (**Extended Data Fig. 5d**), indicating functional recovery. Finally, AZ67 rectified the loss of perinuclear mitochondria observed in induced pluripotent stems cells (iPSC)-derived neural precursor cells (NPC) from CLN7 patients (**Extended Data Fig. 6a-h**). Abnormal accumulation of mitochondria has also been reported in several forms of lysosomal storage diseases^36^, although their functional characterization is missing and the impact on other Batten disease pathogenesis unknown. It would be interesting to ascertain whether the bioenergetic alterations herein described in CLN7 disease are shared with other NCLs. If so, pharmacological inhibition of PFKFB3 would be a suitable therapeutic approach worth testing to delay and/or palliate the devastating consequences of each type of currently intractable^37^ Batten disease.

**Fig. 4.**
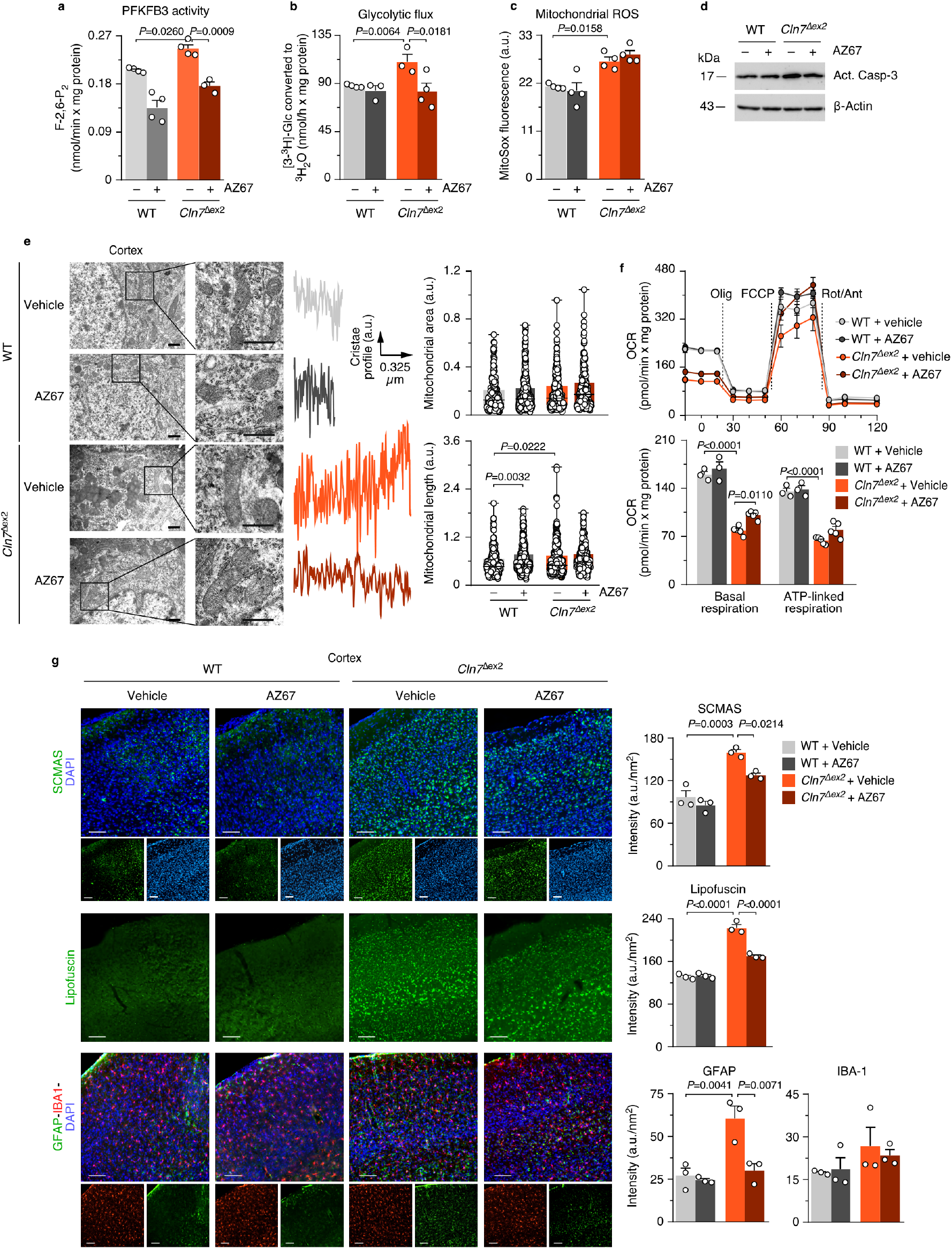
Inhibition of glycolysis by pharmacologically targeting PFKFB3 restores mitochondrial alterations and hallmarks of Cln7^Δex2^ disease *in vivo.* **(a,b,c,d)** Analysis of PFKFB3 activity (**a**), glycolytic flux (**b**), mitochondrial ROS (**c**) and active caspase-3 by western blot (**d**), in primary neurons incubated with the PFKFB3 inhibitor AZ67 (10 nM, 24 h). (**a**, n=3-4; **b**, n= 3-4; **c**, n=3-4; ß-actin, loading control). **(e)** Representative electron microscopy images and analyses of the mouse brain cortex, after 2 months of a daily intracerebroventricular administration of AZ67 (1 nmol/mouse), displaying the cristae profile plot of intensities over the maximal axis of the magnified shown mitochondrion (~350 mitochondria per condition; 3-months old mice). Scale bars, 600 nm. **(f)** OCR analysis (up) and calculated parameters (down) in primary neurons incubated with the PFKFB3 inhibitor AZ67 (10 nM, 24 h) (n=3-5). **(g)** SCMAS, lipofuscin, GFAP and IBA-1 immunohistochemical analysis of the mouse brain cortex after 2 months of a daily intracerebroventricular administration of AZ67 (1 nmol/mouse) (3 serial slices per mouse; n=3 of 3-months-old mice). Scale bar, 100 μm. Data are mean ± S.E.M. for the indicated number of culture preparations or mice. Statistical analyses performed by one-way ANOVA followed by Tukey’s (**a, c, e, f, g, h**) or DMS’s (**b**) post-hoc tests. Representative western blot out of n≥3 experiments is shown (see Extended Data Fig. 4).

## Acknowledgements

We acknowledge the technical assistance of Monica Resch, Monica Carabias-Carrasco, Lucia Martin and Estefania Prieto-Garcia, from the University of Salamanca. This work was funded by the European Union’s Horizon 2020 Research and Innovation Programme (BATCure grant No. 666918 to JPB, SEM, DLM, SS and TRM), MICINN (PID2019-105699RB-I00 / AEI / 10.13039/501100011033 and RED2018 - 102576 - T to JPB; SAF2017-90794-REDT to AA), Instituto de Salud Carlos III (CB16/10/00282 to JPB), Junta de Castilla y León (Escalera de Excelencia CLU-2017-03 to JPN and AA), Ayudas Equipos Investigación Biomedicina 2017 Fundación BBVA (to JPB), Fundación Ramón Areces (to JPB), Instituto de Salud Carlos III (PI18/00285; RD16/0019/0018 to AA), European Regional Development Fund, European Union’s Horizon 2020 Research and Innovation Programme (Grant Agreement 686009 to AA), Junta de Castilla y León (IES007P17 to AA) and Fundación Ramón Areces (to AA). SM benefits from MRC funding to the MRC Laboratory for Molecular Cell Biology University Unit at UCL (award code MC_U12266B) towards lab and office space. Part of this work was funded by Gero Discovery L.L.C. MGM is a ISCIII-Sara Borrel contract recipient (CD18/00203).

## Authors contributions

Conceived the idea: JPB

Designed research: JPB, ILF, AA, POF, SEM, DLM, TM

Performed research: ILF, MGM, CB, OB, NB, BMF, PAB, CVG, DJB, RQC, EF, AS, MGF, LF, CDT

Analyzed data: JPB, ILF, MGM, CB, OB, NB, AA, AS, MGF, LF, TM

Contributed materials: POF, SS

Wrote the manuscript: JPB, ILF, MGM

Edited and approved the manuscript: All co-authors

## Data availability statement

All data generated or analysed during this study are included in this published article (and its supplementary information files).

---

**Extended Data Fig. 1. Related to Fig. 1.**
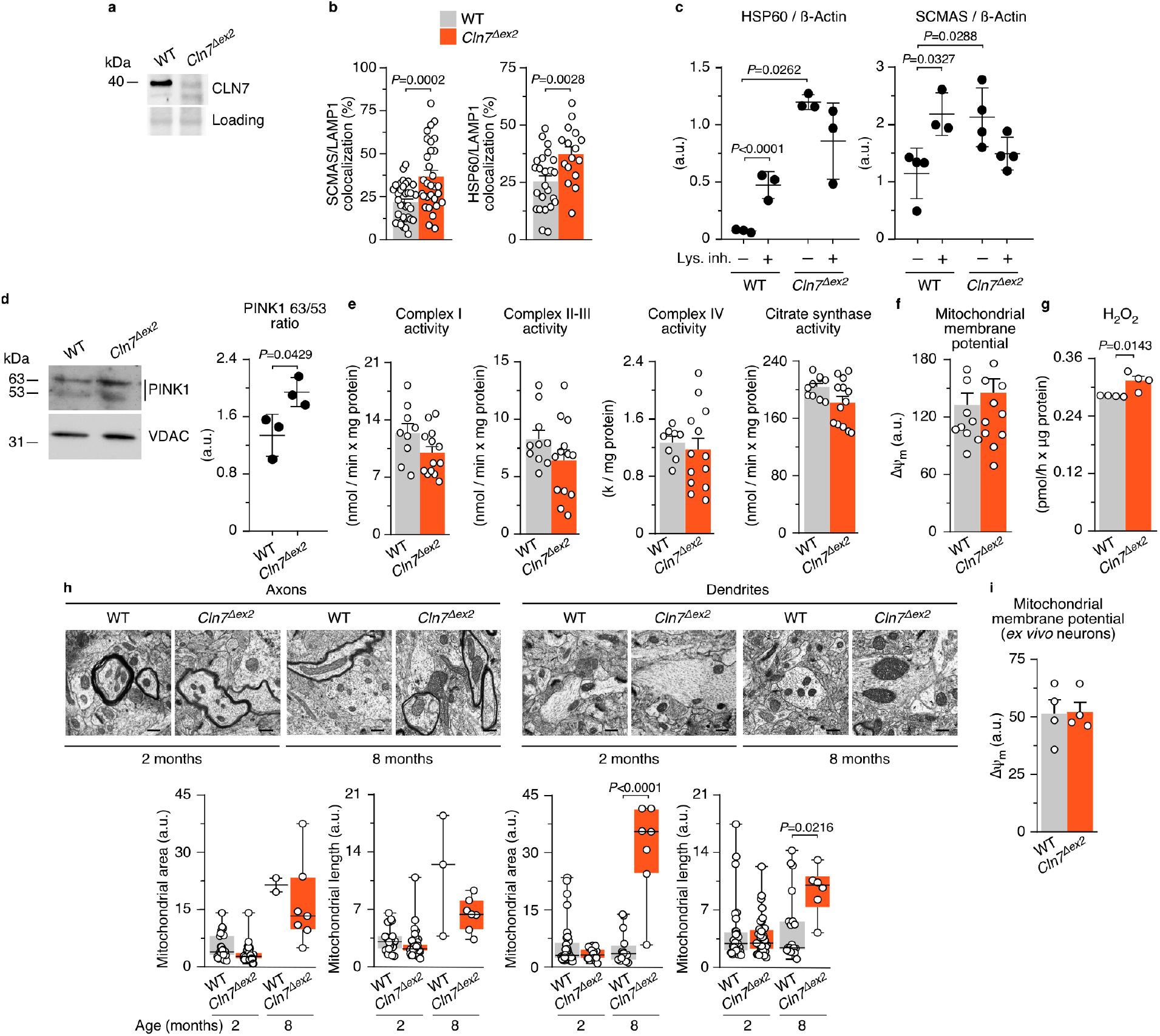
**(a**) Cln7 representative western blot analysis in primary neurons (control loading, Ponceau staining). **(b)** SCMAS/LAMP1 and HSP60/SCMAS co-localization confocal analyses in primary neurons (30 images; n=3) **(c)** Densitometric quantification of the western blot bands shown in Fig. 1b and the replicas in Supplementary Information (n=3-4). **(d)** PINK1 western blot analysis of 63 and 53 kDa bands in primary neuronal mitochondria (VDAC, loading control) (left). Densitometric quantification of the western blot bands shown and the replicas in Supplementary Information (right) (n=4). **(e)** Activities of the mitochondrial respiratory chain complexes I, II-III and IV, and citrate synthase, in cell homogenates from primary neurons (n=9-13). **(f)** Monitorization of Δψ_m_ for MitoSox analysis of Fig. 1e in primary neurons (n=11-13). (**g**) H_2_O_2_ abundance as assessed by AmplexRed fluorescence in primary neurons (n=4). **(h)** Representative electron microscopy images and analyses of mouse brain cortex mitochondria (~40 mitochondria per condition). Scale bar, 500 nm. **(i)** Monitorization of Δψ_m_ for MitoSox analysis of Fig. 1h in freshly isolated neurons from mouse brain cortex (n=4 mice). Data are represented as mean ± S.E.M. (**b,c, d, e, f,g,i**) or box plot (minimum to maximum) (**h**) for the indicated number of culture preparations, mice or mitochondria. Statistical analyses were performed by one-way ANOVA followed by DMS’s post-hoc tests (**c**) or Student’s t test (**d, e, f, g, h,i**).

**Extended Data Fig. 2. Related to Fig. 2.**
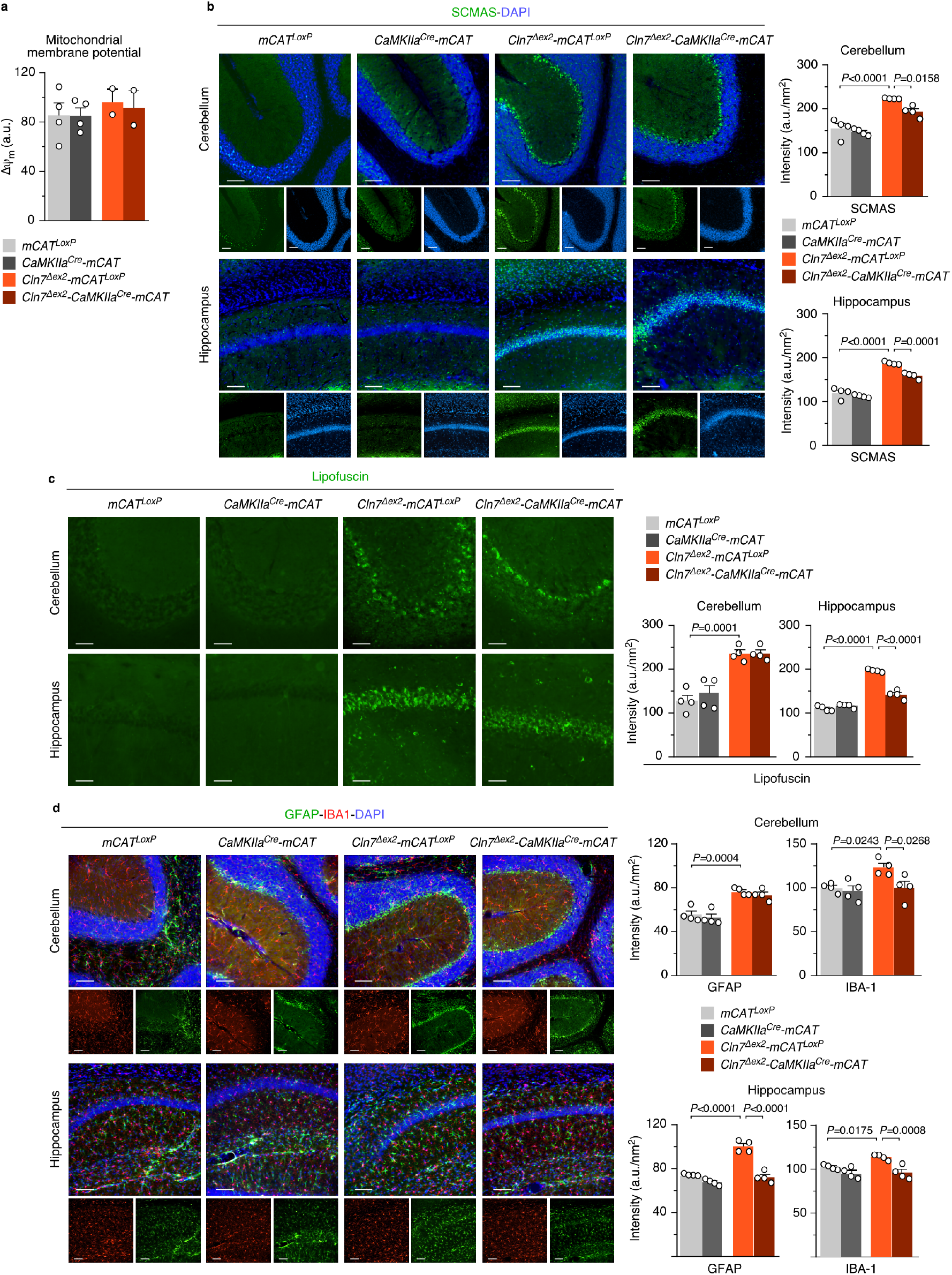
**(a)** Monitorization of Δψ_m_ for MitoSox analysis in Fig. 2a in primary neurons. Data are the mean ± S.E.M. values from 2-4 culture preparations. **(b,c,d)** SCMAS, lipofuscin, GFAP and IBA-1 immunohistochemical analysis of the mouse brain hippocampus and cerebellum. Data are mean ± S.E.M. from 3 serial slices per 3 months-old mouse (**b,d**; n=4 mice; one-way ANOVA followed by post-hoc Tukey), (**c**; n=4 mice; one-way ANOVA followed by DMS’s post-hoc test for cerebellum and Tukey’s post-hoc test for hippocampus). Scale bar, 100 μm.

**Extended Data Fig. 3. Related to Fig. 3.**
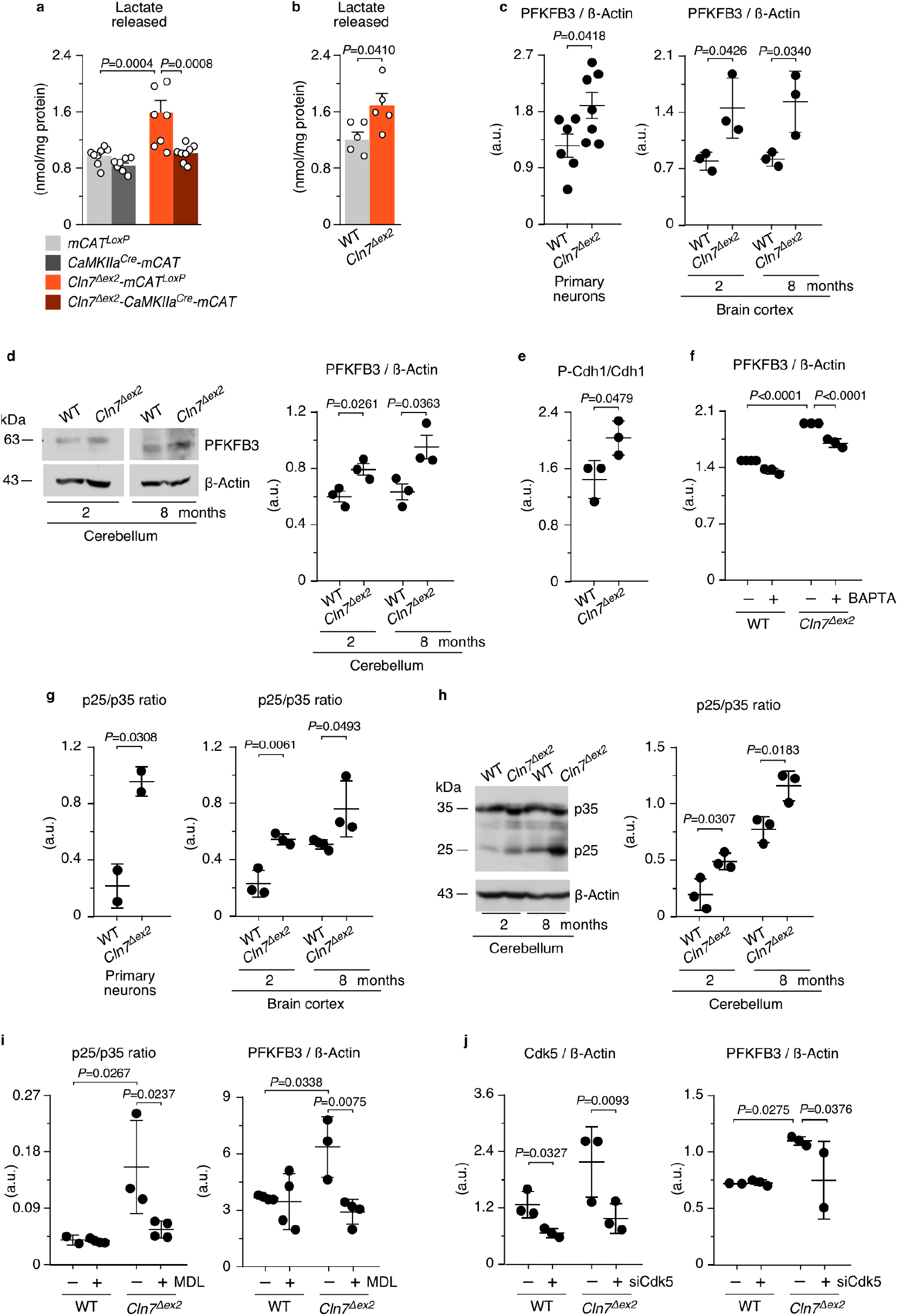
**(a,b)** Lactate release to the culture media in primary neurons. (**a**, n=4-8; **b**; n=5). (**c**) Densitometric quantification of the western blot bands shown in Fig. 3e and replicas in Supplementary Information (n=6-7 mice). (**d**) Representative western blot showing PFKFB3 protein in cerebellum (ß-actin, loading control) (left); densitometric quantification of these bands and replicas in Supplementary Information (right) (n=3 mice). **(e,f,g)** Densitometric quantification of the western blot bands shown in Fig. 3g (**e**), Fig. 3i (**f**) and Fig. 3k (**g**), and replicas in Supplementary Information (n=3). **(h)** Representative western blot showing p35 and its cleavage product p25 in cerebellum (ß-actin, loading control) (left); densitometric quantification of these bands and replicas in Supplementary Information (right) (n=3 mice). **(i,j)** Densitometric quantification of the western blot bands shown in Fig. 3l (**i**), Fig. 3m (**j**), and replicas in Supplementary Information (**i**, n=3-4; **j**, n=2-3). Data are mean ± S.E.M. (**a, b**) or S.D. (**c-j**) for the indicated number of culture preparations or mice. Statistical analyses were performed by one-way ANOVA followed by Tukey’s (**a,c,d,f,i**) or DMS’s (**j**) post-hoc tests, or Student’s t test (**b,e,g,h**).

**Extended Data Fig. 4. Related to Fig. 4.**
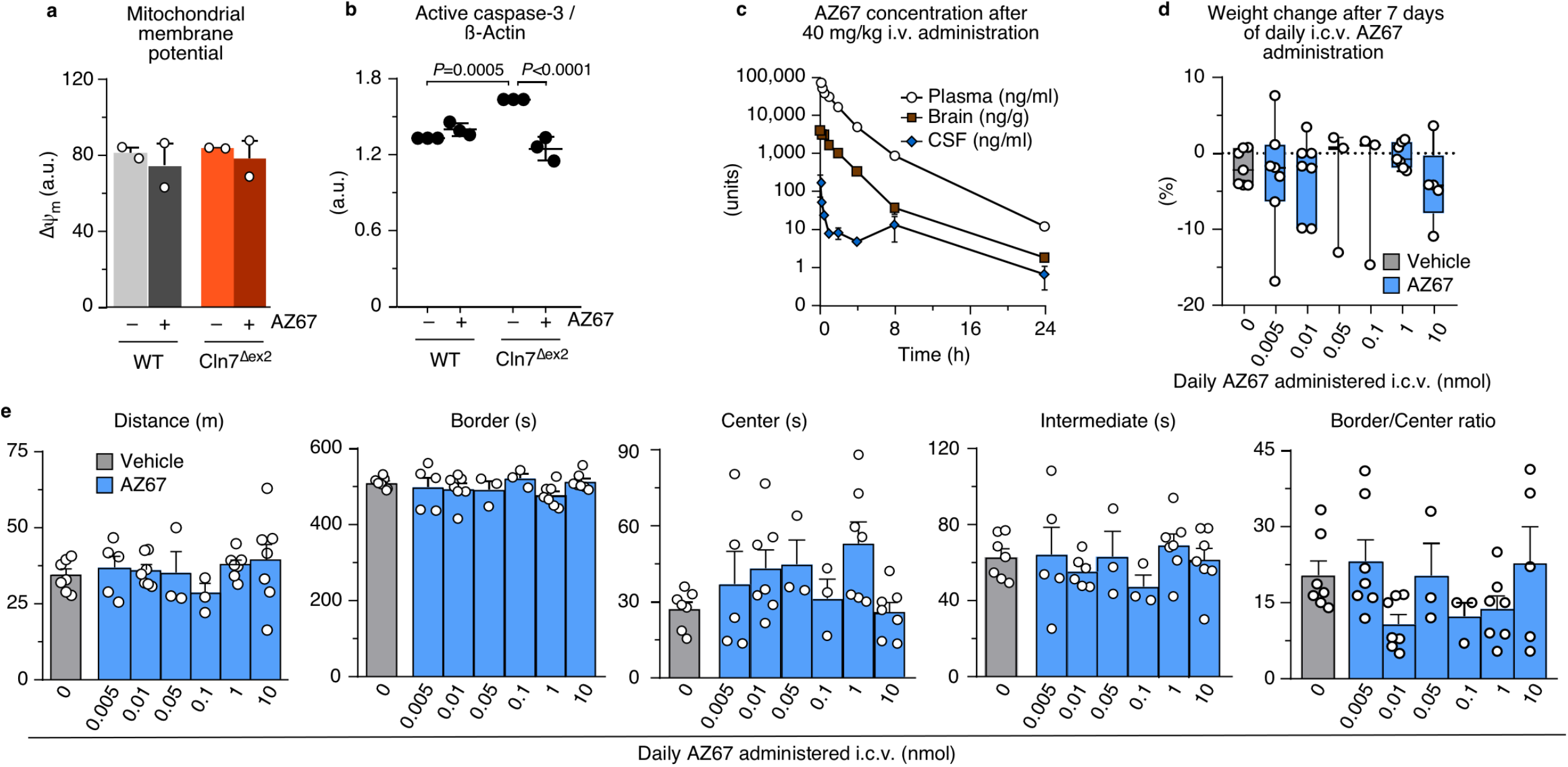
(**a**) Monitorization of Δψ_m_ for MitoSox analysis of Fig. 4c in primary neurons. Data are mean ± S.E.M. (n= 7-8 culture preparations). **(b)** Densitometric quantification of the western blot bands shown in Fig. 4d and replicas in Supplementary Information. Data are mean ± S.D. (n=3 culture preparations; oneway ANOVA followed by Tukey’s post-hoc test). **(c)** Time course evaluation of AZ67 concentration in plasma, brain and cerebrospinal fluid after the intravenous administration of 40 mg/kg of AZ67 in C57Bl/6 mice. Data are mean ± S.E.M. (n=3 mice). Related to Fig. 4e,g. **(d)** Dose response evaluation of the weight fluctuations after 7 days of a daily intracerebroventricular injection of AZ67 (1 nmol/mouse). Data are represented in box plots (minimum to maximum) (n=3-7 mice). Related to Fig. 4e,g. **(e)** Dose response evaluation of the open field test after 7 days of a daily intracerebroventricular injection of AZ67 (1 nmol/mouse). Total distance, time spend in border, center and intermediate zone, and the ratio of border to center time are represented. Data are mean ± S.E.M. (n=3-7 mice). Related to Fig. 4e,g.

**Extended Data Fig. 5. Related to Fig. 4.**
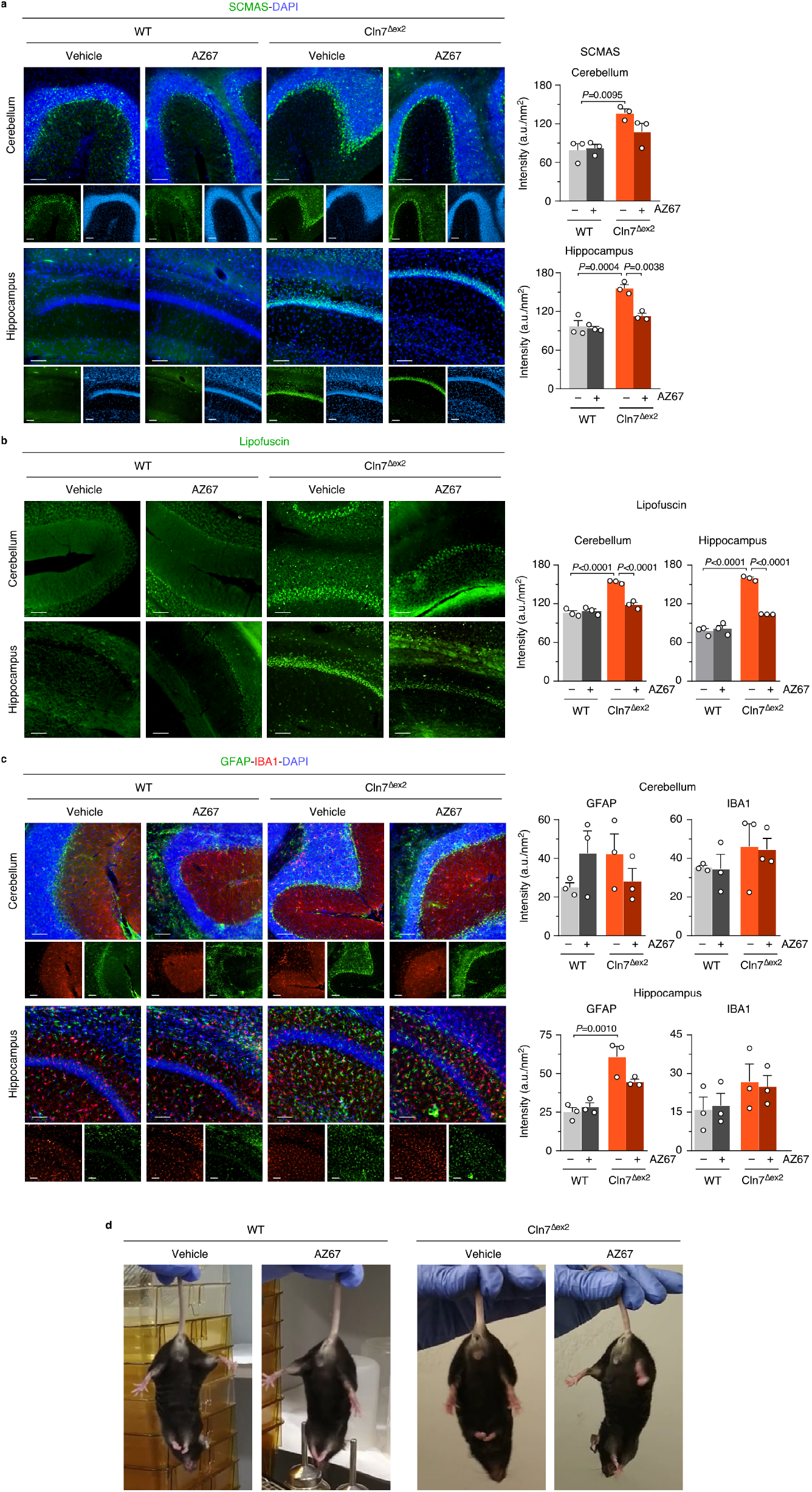
**(a,b,c)** Immunohistochemical analysis of the cerebellum and hippocampus of 3 months-old mice after 2 months of a daily intracerebroventricular administration of AZ67 (1 nmol/mouse) against SCMAS (**a**), lipofuscin (autofluorescence) (**b**), GFAP and IBA-1 (**c**). Data are mean ± S.E.M. from the quantification of 3 serial slices per mouse in 3 mice (one-way ANOVA followed by Tukey’s post-hoc test). Scale bar, 100 μm. Related to Fig. 4g. **(d)** Representative pictures of 3 months-old mice after 2 months of a daily intracerebroventricular administration of AZ67 (1 nmol/mouse), or vehicle, showing hindlimb paralysis phenotype in the *Cln7*^Δ*ex2*^ mice. See supplementary videos.

**Extended Data Fig. 6. Related to Fig. 4.**
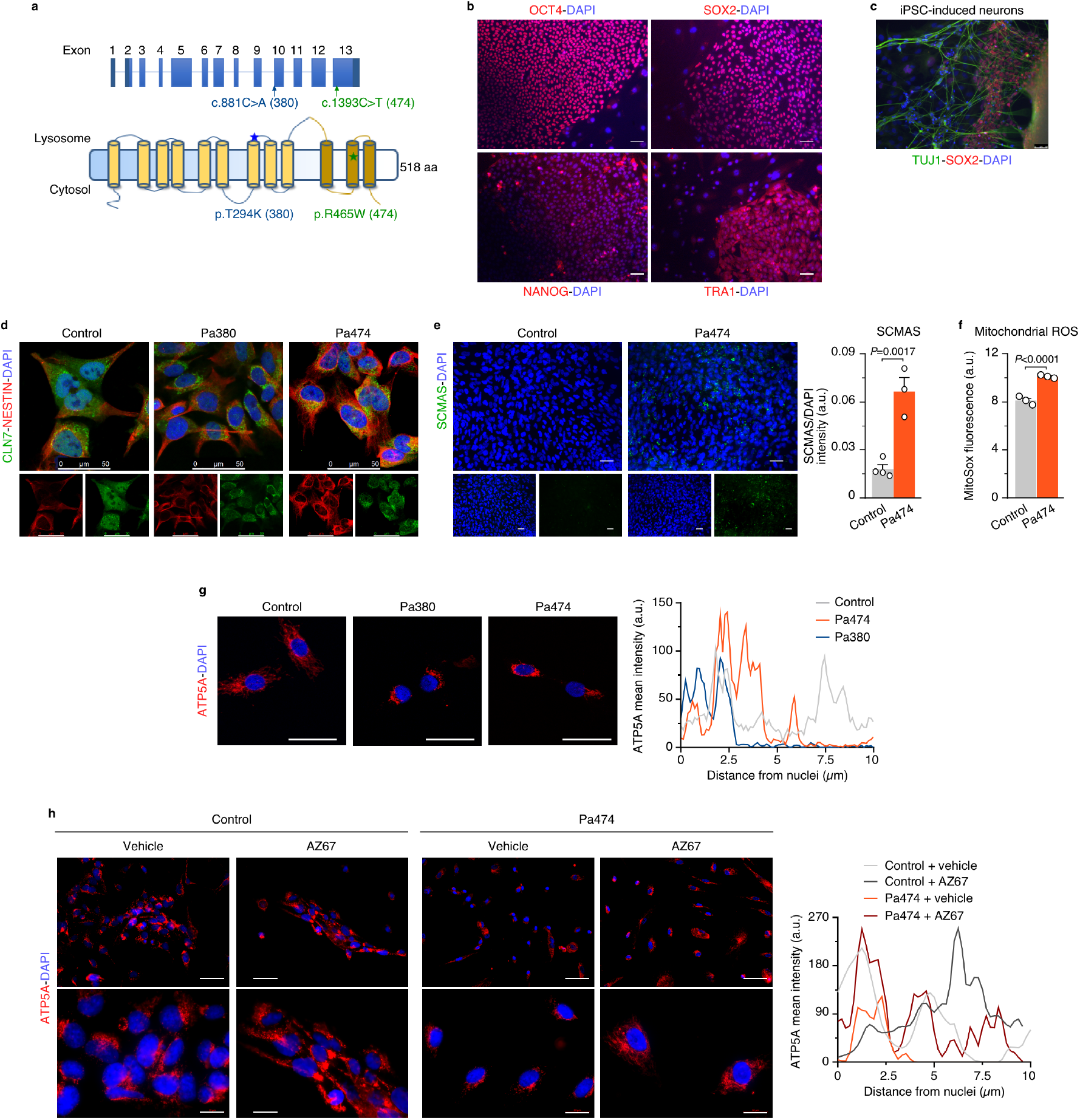
**(a)** Schematic representation of the locations of the CLN7 mutations found in patient 380 (Pa380, c.881C>A; pT294K) and patient 474 (Pa474, c.1393C>T; p.R465W). **(b)** iPSC characterization in Pa474 with the pluripotency markers OCT4, SOX2, Nanog and Tra-1-60. Scale bar, 50 μm. **(c)** Characterization of differentiated neurons derived from iPSC in Pa474. Scale bar, 50 μm. **(d)** NPCs characterization in Pa380, Pa474 and a healthy, age-matched control individual. Scale bar, 50 μm. **(e)** Immunocytochemical analysis of SCMAS abundance in NPCs derived from Pa474 iPSC. Data are the mean ± S.E.M. values from three independent culture preparations (Student’s t test). Scale bar, 50 μm. **(f)** Mitochondrial ROS analysis in NPCs. Data are the mean ± S.E.M. values from three independent culture preparations (Student’s *t* test). **(g)** Immunocytochemical analysis of mitochondrial marker ATP5A in NPCs derived from Pa380, Pa474 and healthy-matched control patient. Scale bar, 50 μm. The right panel shows a representative pixel intensity profile of ATP5A across the maximal axis of the cell that departs from the nucleus. **(h)** NPCs derived from Pa474 iPSC were incubated with AZ67 for 24h, fixed and subjected to immunocytochemical analysis for ATP5A. Scale bars, 60 μm (upper images) and 20 μm (lower images). The right panel shows a representative pixel intensity profile of ATP5A across the maximal axis of the cell that departs from the nucleus.

## Online Methods

### Animals

All protocols were performed according to the European Union Directive 86/609/EEC and Recommendation 2007/526/EC, regarding the protection of animals used for experimental and other scientific purposes, enforced in Spanish legislation under the law 6/2013. All protocols were approved by the Bioethics Committee of the University of Salamanca. Animals were bred at the Animal Experimentation Facility of the University of Salamanca in cages (maximum of five animals per cage), and a light–dark cycle was maintained for 12h. The humidity was 45–65%, and the temperature was 20–25 °C. Animals were fed *ad libitum* with a standard solid diet (17% proteins, 3% lipids, 58.7% carbohydrates, 4.3% cellulose, 5% minerals and 12% humidity) and given free access to water. Cln7 knockout mouse carrying the European Conditional Mouse Mutagenesis (EUCOMM) tm1d allele by Cre-mediated recombination of the floxed exon 2 of the murine Cln7/Mfsd8 gene (Cln7^Δex2^)^11^. To abrogate mitochondrial ROS selectively in neurons in the Cln7^Δex2^ mice *in vivo*, we crossed Cln7^Δex2^ with transgenic mice harbouring the full-length cDNA encoding catalase fused to the cytochrome *c* oxidase subunit VIII–mitochondrial leading sequence (mitoCatalase or mCAT), which has incorporated a floxed transcriptional STOP cassette between the mitochondrial-tagged catalase cDNA and the CAG promoter, which were previously generated in our laboratory by homologous recombination in the *Rosa26* locus under a C57BL/6 background (mCAT^LoxP^/+) in order to achieve tissue- and time-specific expression of mCAT *in vivo*^18^. mCAT^LoxP^/+ mice were mated with mice harbouring Cre recombinase under control of the neuronal specific CAMKII promoter (CAMKIIa^Cre^). The progeny, namely CAMKIIa^Cre^/+; mCAT^LoxP^/+, were crossed with Cln7^Δex2^/Cln7^Δex2^ mice^11^. The offspring were crossed to obtain the following littermates genotypes: i) +/+; mCAT/+; +/+ (mCAT^LoxP^); ii) +/+; mCAT/+; CamKIIa/+ (CAMKIIa^Cre^-mCAT); iii) Cln7^Δex2^/Cln7^Δex2^; mCAT/+; +/+ (Cln7^Δex2^-mCAT^LoxP^); iv) Cln7^Δex2^/Cln7^Δex2^; mCAT/+; CamKIIa/+ (Cln7^Δex2^-CAMKIIa^Cre^-mCAT).

### Genotyping by polymerase chain reaction (PCR)

For Cln7^Δex2^ genotyping, a PCR with the following primers was performed 5’-TGGTGCATTAATACAGTCCTAGAATCCAGG-3’, 5’-CTAGGGAGGTTCAGATAGTAGAACCC-3’, 5’-TTCCACCTAGAGAATGGAGCGAGATAG-3’, resulting in a 290 bp band in the case of Cln7^Δex2^ mice, and 400 bp for wild type^11^. The primer sequences for genotyping the mCAT^LoxP^ allele were 5’-CTCCCAAAGTCGCTCTGAGTTGTTATCA-3’, 5’-CGATTTGTGGTGTATGTAACTAATCTGTCTGG-3’ and 5’-GCAGTGAGAAGAGTACCACCATGAGTCC-3’, which yielded a 778-bp band for the wild-type allele and a 245-bp band for the mCAT^LoxP^ allele^18^. CaMKIIa^Cre^ transgene was detected by amplifying a 270 bp region of Cre recombinase by PCR. Forward and reverse oligonucleotides used were, respectively, 5’-GCATTTCTGGGGATTGCTTA-3’ and 5’-CCCGGCAAAACAGGTAGTTA-3’. An internal control was used to detect false negatives using the endogenous SCNA gene. Its forward and reverse oligonucleotides were, respectively, 5’-ATCTGGTCCTTCTTGACAAAGC-3’ and 5’-AGAAGACCAAAGAGCAAGTGACA-3’, which generated a 150 bp band.

### Reverse transcription-real time quantitative PCR (RT-qPCR)

This was performed in total RNA samples, purified from primary culture of neurons using the GenElute Mammalian Total RNA Miniprep Kit (Sigma), following the manufacturer’s protocol. Amplifications were performed in 100 ng of RNA, using Power SYBR Green RNA-to-CT 1-Step kit (Applied Biosystems). The primers were 5’-TTCTCAGGTTTTTGCGGAGAAC-3’ and 5’-GTGCACATGTATGAGCTGGCA-3’ for PFKFB3 and 5’-CGATGCCCTGAGGCTCTTTT-3’ and 5’-CAACGTCACACTTCATGATG-3’ for β-actin. The mRNA abundance of each transcript was normalized to the β-actin mRNA abundance obtained in the same sample. WT neurons were used as a control.

### *In vitro* Cre recombinase activity induction

Infection with adenovirus carriers of Cre recombinase and empty adenovirus (Control) was used to induce mCAT expression in primary culture of neurons conditionals for mCAT expression (mCat^LoxP^ and Cln7^Δex2^-mCAT^LoxP^). Virus, transduced at 10 MOI, were purchased to Gene Transfer Vector Core (University of Iowa). Transduction was performed 3 days before cell recollection, and viral particles were left in the cultures during 24h.

### Primary cultures

Primary cultures of mice cortical neurons were prepared from the offspring of 14.5 days Cln7^Δex2^ pregnant^11^ or WT pregnant (C57BL/6J) mice according to a previously validated protocol^38^. In essence, cells were seeded at 2.0·10^5^ cells per cm^2^ in different-sized plastic plates coated with poly-D-lysine (10 μg/mL) and incubated in Neurobasal-A (Life Technologies) supplemented with 5.5 mM of glucose, 0.25 mM of sodium pyruvate, 2 mM glutamine and 2% (vol/vol) B27 supplement (Life Technologies). At 72 h after plating, medium was replaced, and cells were used at day 7. Cells were incubated at 37 °C in a humidified 5% (vol/vol) CO_2_-containing atmosphere.

### Induced pluripotent stem cells (iPSC) and Neural Progenitor Cells (NPC) generation

iPSC were generated from two CLN7 patients (Pa380 and Pa474), then characterized and differentiated to NPC as previously described^39^. Human iPSC-derived NPCs from a control patient and patients Pa380 (c.881C>A; pT294K) and Pa474 (c.1393C>T; p.R465W) harboring the indicated CLN7 homozygous mutations, were plated on Matrigel® Matrix in Nunc™ Lab-Tek™ 8-well Chamber Slides and cultured in Neural Expansion Medium (NEM) with DMEM/F12, NEAA, N-2 supplement, B-27 supplement, heparin, bFGF protein, penicillin/streptomycin. iPSCs pluripotency was confirmed by immunocytochemistry using OCT4 (1:200, ab19857; Abcam), SOX2 (1:100, AF2018; R&D Systems), Nanog (1:100, ab21624: Abcam) and Tra-1-60 (1:200, MAB1295; R&D Systems), and by confirming their ability to differentiate into neurons using TUJ-1 (1:200, MAB1195; R&D Systems) staining. NPC identity was confirmed by Nestin^+^/SOX2^-^ immunostaining.

### Freshly purification of neurons from the brain cortex form adult mice

Adult mouse brain (from 6 months-old animals) tissue was dissociated with the Adult Brain Isolation Kit (Miltenyi). Dissociated cells, after removal of debris and red blood cells, neurons were separated with the Neuron Isolation Kit (Miltenyi), accordingly to the manufacturer’s protocol. The identity of the isolated fraction was confirmed previously^16^ by western blot against the neuronal marker microtubule associated protein 2 (MAP2).

### Cell treatments

Neurons in primary culture were incubated with the rationally-designed, potent and highly selective PFKFB3 inhibitor AZ PFKFB3 67 (herein referred as AZ67)^34^ (Tocris) (10 nM) or the calpain inhibitor MDL-28170 (MDL, 10 μM; Sigma) for 24 h. The cell permeable Ca^2+^-quelator BAPTA was used in primary culture of neurons in the presence of Hanks’s solution without calcium (134.2 mM NaCl, 5.26 mM KCl, 0.43 mM KH_2_PO_4_, 4.09 mM NaHCO_3_, 0.33 mM Na_2_HPO_4_·2H_2_O, 5.44 mM glucose, 20 mM Hepes, pH 7.4) for 1 h (10 μM; Sigma). NPCs were incubated with AZ67 (10 nM) for 24 h in NEM.

### Autophagy-lysosomal pathway measurement

To analyze the autophagy-lysosomal pathway, primary neurons were incubated in the absence or presence of the inhibitors of the lysosomal proteolysis leupeptin (100 μM) and ammonium chloride (20 mM) for 1 h. Cells were lysed and immunoblotted against SCMAS and HSP60 to assess mitophagic flux^13^.

### Cell transfections

For knockdown experiments, small interfering RNA (siRNA) against CDK5 (siCDK5) (s201147; Thermo Fisher) was used. An siRNA control (siControl) (4390843; Thermo Fisher) was used in parallel. siRNA transfections were performed using the Lipofectamine RNAiMAX reagent (Thermo Fisher) according to the manufacturer’s protocol at an siRNA final concentration of 9 nM. Cells were used after 3 days.

### Total membrane purification

Membranes were isolated from primary cultures of neurons, or whole brain homogenates, following a previously reported protocol^11^. Briefly cells or tissue were homogenized in sucrose buffer (200 mM sucrose, 50 mM Tris-HCl; pH 7.5, 1 mM EDTA) and centrifuged 5 minutes at 1,500 g. Supernatants were centrifuged at 20,000 g 10 minutes. Pellet was then homogenized in extraction buffer (50 mM Tris-HCl; pH 7.5, 1% (vol/vol) Triton-X100, 1 mM EDTA) and incubated 30 min on ice, followed by a centrifugation at 20,000 g 10 minutes. Membranes enriched fraction remained in the supernatant.

### Western Blotting

Cells were lysed in RIPA buffer (1% sodium dodecylsulfate, 10 mM ethylenediaminetetraacetic acid (EDTA), 1 % (vol/vol) Triton X-100, 150 mM NaCl and 10 mM Na2HPO4, pH 7.0), supplemented with protease inhibitor mixture (Sigma), 100 μM phenylmethylsulfonyl fluoride, and phosphatase inhibitors (1 mM o-vanadate). Samples were boiled for 5 min. Aliquots of cell lysates (40 μg of protein) were subjected to SDS/PAGE on an 8 to 12% (vol/vol) acrylamide gel (MiniProtean; Bio-Rad) including PageRuler Prestained Protein Ladder (Thermo). The resolved proteins were transferred electrophoretically to nitrocellulose membranes (0.2 μm, BioRad). Membranes were blocked with 5% (wt/vol) low-fat milk in TTBS (20 mM Tris, 150 mM NaCl, and 0.1% (vol/vol) Tween 20, pH 7.5) for 1 h. Subsequent to blocking, membranes were immunoblotted with primary antibodies overnight at 4 °C. After incubation with horseradish peroxidase conjugated goat anti-rabbit IgG (1/10,000, Santa Cruz Biotechnologies), goat anti-mouse IgG (1/10,000, Bio-Rad), rabbit anti-goat IgG (1/10,000, Abcam) or goat anti-rabbit IgG (1/3,000, Bio-Rad), membranes were immediately incubated with the enhanced chemiluminescence kit WesternBright ECL (Advansta), or SuperSignal West Femto (Thermo) before exposure to Fuji Medical X-Ray film (Fujifilm), and the autoradiograms were scanned. Ponceau staining (Sigma) was occasionally used as an indicator of loading. At least two biologically independent replicates were always performed, although only one representative Western blot is shown in the main figures. The protein abundances of all Western blots per condition were measured by densitometry of the bands on the films using ImageJ 1.48u4 software (National Institutes of Health) and were normalized per the loading control protein. The resulting values were used for the statistical analysis.

### Primary antibodies for western blotting

Immunoblotting was performed with anti-ATP-C (SCMAs) (1/1000) (ab181243; Abcam), anti-VDAC (1/666) (PC548; Calbiochem), anti-HSP60 (1/666) (ab46798; Abcam), anti-PINK1 (1/500) (sc-33796; Santa Cruz Biotechnology), anti-NDUFS1 (1/500) (sc-50132; Santa Cruz Biotechnology), anti-PFKFB3 (1/500) (H00005209-M08; Novus Biologicals), antip25/35 (1/666) (2680; Cell Signalling), anti-caspase-3 (1/2,000) (9661S; Cell Signalling), anti-CLN7 (1/500) (donated by Dr. Stephan Storch) and anti-β-Actin (1/30,000) (A5441; Sigma).

### Immunocytochemistry

Cells were seeded on coverslips, fixed with a 4% paraformaldehyde (PFA) solution, blocked and incubated with primary antibodies overnight at 4°C. The primary antibodies were anti-C-subunit of ATP synthase (SCMAS) (1/200) (ab181243; Abcam), anti heat-shock protein-60 (HSP60) (1/500((ab46798) and anti-LAMP1 (1/100) (1D4B; Developmental Studies Hybridoma Bank). They were then incubated for 1 h with fluorescent secondary antibodies anti-rabbit Alexa Fluor 488 (A-11008; Thermo Fisher), and anti-rat Alexa Fluor 647 (A-21247; Thermo Fisher). DAPI (4’,6-diamidino-2-phenylindole) was used for nuclei visualization. Coverslips were mounted in ProLong Gold antifade reagent. Negative controls were performed with either no primary or no secondary antibodies. No staining was detected in any case. Images were acquired on an Operetta CLS high-content imaging system (PerkinElmer) using 63x (1.15 numerical aperture) objective. Images were acquired at the same exposure times in the same imaging session. Image quantification was performed after appropriate thresholding using the ImageJ software (NIH). The percentage of colocalization was calculated using the JAva COnstraint Programming (JACoP) plugin, specifically Manders’ Overlap Coefficient, in single Z-stack sections.

### Mitochondrial isolation

Mitochondria were obtained according to a previously published protocol^16^. Briefly, cell pellets (10 million) or brain cortex were frozen at −80 °C and homogenized (ten-twelve strokes) in a glass-Teflon Potter–Elvehjem homogenizer in buffer A (83 mM sucrose and 10 mM MOPS; pH 7.2). The same volume of buffer B (250 mM sucrose and 30 mM MOPS) was added to the sample, and the homogenate was centrifuged (1,000 g, 5 min) to remove unbroken cells and nuclei. Centrifugation of the supernatant was then performed (12,000 g, 3 min) to obtain the mitochondrial fraction, which was washed in buffer C (320 mM sucrose; 1 mM EDTA and 10 mM Tris-HCl; pH 7.4). Mitochondria were suspended in buffer D (1 M 6-aminohexanoic acid and 50 mM Bis-Tris-HCl, pH 7.0).

### Blue Native Gel Electrophoresis and in gel activity for complex I

For the assessment of complex I organization, digitonin-solubilized (4 g/g) mitochondria (10–50 μg) were loaded in NativePAGE Novex 3–12% (vol/vol) gels (Life Technologies). After electrophoresis, in-gel NADH dehydrogenase activity was evaluated^16^, allowing the identification of individual complex I and complex I-containing supercomplexes bands. Furthermore, a direct electrotransfer was performed followed by immunoblotting against mitochondrial complex I antibody NDUFS1. Direct transfer of BNGE was performed after soaking the gels for 20 min (4 °C) in carbonate buffer (10 mM NaHCO_3_; 3 mM Na_2_CO_3_·10H_2_O; pH 9.5–10). Proteins transfer to polyvinylidene fluoride (PVDF) membranes was carried out at 300 mA, 60 V, 1 h at 4 °C in carbonate buffer.

### Determination of PPP and glycolytic fluxes

These were measured in 8 cm^2^ flasks of primary cultures of neurons containing a central microcentrifuge tube with either 0.8 ml benzethonium hydroxide (Sigma) for ^14^CO_2_ equilibration or 1 ml H_2_O for ^3^H_2_O equilibration. Incubations were carried out in KRPG (NaCl 145 mM; Na2HPO4 5.7 mM; KCl 4.86 mM; CaCl2 0.54 mM; MgSO4 1.22 mM; pH 7.35) containing 5 mM D-glucose at 37 °C in the air-thermostatized chamber of an orbital shaker. To ensure adequate oxygen supply for oxidative metabolism throughout the incubation period, flasks were filled with oxygen (5% CO_2_/O_2_) before being sealed. To measure the carbon flux from glucose through the PPP, cells were incubated in KRPG (5 mM D-glucose) buffer supplemented with 0.5 μCi d-[1-^14^C]glucose or [6-^14^C]glucose for 90 min, as previously described^21,40^. Incubations were terminated by the addition of 0.2 ml 20% perchloric acid (Merck Millipore), and 40 min before the benzethonium hydroxide (containing ^14^CO_2_) was removed, and the radioactivity was measured with a liquid scintillation analyzer (Tri-Carb 4810 TR, PerkinElmer). PPP flux was calculated as the difference between ^14^CO_2_ production from [1-^14^C]glucose (which decarboxylates through the 6-phosphogluconate dehydrogenase-catalyzed reaction) and that of [6-^14^C]glucose (which decarboxylates through the TCA cycle)^21,41^. Glycolytic flux was measured by assaying the rate of ^3^H_2_O production from [3-^3^H]glucose through a similar method, but incubating cells with 3 μCi d-[3-^3^H]glucose in KRPG buffer per flask for 120 min, as previously described^21,40^. Incubations were terminated with 0.2 ml 20% perchloric acid, and the cells were further incubated for 96 h to allow for ^3^H_2_O equilibration with H_2_0 present in the central microcentrifuge tube. The ^3^H_2_O was then measured by liquid scintillation counting (Tri-Carb 4810 TR, PerkinElmer). Under these experimental conditions, 75% of the produced ^14^CO_2_ or 28% of the produced ^3^H_2_O was recovered and used for the calculations as previously established^40^.

### Lactate determination

Lactate concentrations were measured in the culture medium espectrophotometrically^21^ by determination of the increments in the absorbance of the samples at 340 nm in a mixture containing 1 mM NAD^+^, 8.25 U lactate dehydrogenase in 0.25 M glycine, 0.5 M hydrazine and 1 mM EDTA buffer, pH 9.5.

### Fructose-2,6-bisphosphate determinations

For F-2,6-P2 determinations, cells were lysed in 0.1 M NaOH and centrifuged (20,000 g, 20 min). An aliquot of the homogenate was used for protein determination, and the remaining sample was heated at 80 °C (5 min), centrifuged (20,000 g, 20 min) and the resulting supernatant used for the determination of F-2,6-P2 concentrations using the coupled enzymatic reaction as previously described^42^.

### Phos-tag SDS-PAGE

For the evaluation of phosphorylation levels of CDH1, primary cultures of neurons were homogenized in extraction buffer (100 mM NaCl, 50 mM Tris pH 8, 1% (vol/vol) NP40). Electrophoresis was performed in 8 % (vol/vol) SDS-PAGE gels in the presence of 37.5 μM of PhosTag Acrylamide (ALL-107M, Wako) and 75 μM of MnCl2. After electrophoresis, gels were washed 3 times in transfer buffer with 1mM of EDTA, before electroblotting.

### Mitochondrial ROS

Mitochondrial ROS were determined with the fluorescent probe MitoSox (Life Technologies). Neurons, from primary cultures or adult brain-cell suspensions, were incubated with 2 μM of MitoSox for 30 min at 37 °C in a 5% CO_2_ atmosphere in HBSS buffer (134.2 mM NaCl, 5.26 mM KCl, 0.43 mM KH_2_PO_4_, 4.09 mM NaHCO_3_, 0.33 mM Na_2_HPO_4_·2H_2_O, 5.44 mM glucose, 20 mM HEPES and 20 mM CaCl_2_·2H_2_O, pH 7.4). The cells were then washed with phosphate-buffered saline (PBS: 136 mM NaCl; 2.7 mM KCl; 7.8 mM Na_2_HPO_4_·2H_2_O; 1.7 mM KH_2_PO_4_; pH 7.4) and collected by trypsinization. MitoSox fluorescence intensity was assessed by flow cytometry (FACScalibur flow cytometer, BD Biosciences) and expressed in arbitrary units.

### H_2_O_2_ Determination

For H_2_O_2_ assessments, AmplexRed (Life Technologies) was used. Cells were tripsinized and incubated in KRPG buffer (145 mM NaCl, 5.7 mM Na_2_HPO_4_, 4.86 mM KCl, 0.54 mM CaCl_2_, 1.22 mM MgSO4, 5.5 mM glucose, pH 7.35) in the presence of 9.45 μM AmplexRed containing 0.1 U/mL horseradish peroxidase. Luminescence was recorded for 2 h at 30 min intervals using a Varioskan Flash (Thermo Scientific) (excitation, 538 nm; emission, 604 nm). Slopes were used for calculations of the rates of H_2_O_2_ formation.

### Mitochondrial membrane potential

The mitochondrial membrane potential (ΔΨ_m_) was assessed with MitoProbe DiIC1(5) (Life Technologies) (50 nM) by flow cytometry (FACScalibur flow cytometer, BD Biosciences) and expressed in arbitrary units. For this purpose, cell suspensions were incubated with the probe 30 min at 37°C in PBS. ΔΨ_m_ are obtained after subtraction of the potential value determined in the presence of carbonyl cyanide-4-(trifluoromethoxy)phenylhydrazone (FCCP) (10 μM, 15 min) for each sample.

### Cytosolic Ca^2+^ determination using Fura-2 fluorescence

To estimate the intracellular Ca^2+^ levels in neurons we used the fluorescent probe Fura-2 (acetoxymethyl-derivative; Life Technologies), as previously described^43^. Essentially, neurons were incubated with Fura-2 (2 *μ*M) for 40 min in Neurobasal medium at 37 °C. Then, cells were washed and further incubated with standard buffer (140 mM NaCl, 2.5 mM KCl, 15 mM Tris-HCl, 5 mM D-glucose, 1.2 mM Na_2_HPO_4_, 1 mM MgSO4 and 1 mM CaCl_2_, pH 7.4) for 30 min and 37 °C. Finally, the standard buffer was removed and experimental buffer (140 mM NaCl, 2.5 mM KCl, 15 mM Tris-HCl, D-glucose, 1.2 mM Na_2_HPO_4_, and 2 mM CaCl_2_, pH 7.4) was added. Emissions at 510 nm, after excitations at 335 and 363 nm, respectively, were recorded at in a Varioskan Flash (Thermo) spectrofluorometer at 37 °C. Ca^2+^ levels were estimated by representing the ratio of fluorescence emitted at 510 nm obtained after excitation at 335 nm divided by that at 363 nm (F335/F363). Background subtraction was accomplished from emission values obtained in Fura-2-lacking neurons. At least, 6 wells were recorded per condition in each experiment and the averaged values are shown, normalized per mg of protein present in the sample.

### Bioenergetics

Oxygen consumption rates of neurons were measured in real-time in an XFe24 Extracellular Flux Analyzer (Seahorse Bioscience). The instrument measures the extracellular flux changes of oxygen in the medium surrounding the cells seeded in XFe24-well plates. Assay was performed on the day 7 after cell plating/culture. Regular cell medium was then removed and cells were washed twice with DMEM running medium (XF assay modified supplemented with 5 mM glucose, 2 mM L-glutamine, 1 mM sodium pyruvate, 5 mM HEPES, pH 7.4) and incubated at 37°C without CO_2_ for 30 minutes to allow cells to pre-equilibrate with the assay medium. Oligomycin, FCCP or antimycin/rotenone diluted in DMEM running medium were loaded into port-A, port-B or port-C, respectively. Final concentrations in XFe24 cell culture microplates were 1 μM oligomycin, 2 μM FCCP and 2.5 μM antimycin and 1.25 μM rotenone. The sequence of measurements was as follow unless otherwise described. Basal level of oxygen consumption rate (OCR) was measured 3 times, and then port-A was injected and mixed for 3 minutes, after OCR was measured 3 times for 3 minutes. Same protocol with port-B and port-C. OCR was measured after each injection to determine mitochondrial or non-mitochondrial contribution to OCR. All measurements were normalized to average three measurements of the basal (starting) level of cellular OCR of each well. Each sample was measured in 3–5 wells. Experiments were repeated 3–5 times with different cell preps. Non-mitochondrial OCR was determined by OCR after antimycin/rotenone injection. Maximal respiration was determined by maximum OCR rate after FCCP injection minus non-mitochondrial OCR. ATP production was determined by the last OCR measurement before oligomycin injection minus the minimum OCR measurement after oligomycin injection.

### Activity of Mitochondrial Complexes

Cells were collected and suspended in PBS (pH 7.0). After three cycles of freeze/thawing, to ensure cellular disruption, complex I, complex II, complex II–III, complex IV, and citrate synthase activities were determined. Rotenone-sensitive NADH-ubiquinone oxidoreductase activity (complex I)^44^ was measured in KH_2_PO_4_ (20 mM; pH 7.2) in the presence of 8 mM MgCl2, 2.5 mg/mL BSA, 0.15 mM NADH, and 1 mM KCN. Changes in absorbance at 340 nm (30 °C) (e = 6.81 mM^-1^·cm^-1^) were recorded after the addition of 50 μM ubiquinone and 10 μM rotenone. Complex II-III (succinate–cytochrome c oxidoreductase) activity^45^ was determined in the presence of 100 mM phosphate buffer, plus 0.6 mM EDTA(K^+^), 2 mM KCN, and 200 μM cytochrome *c.* Changes in absorbance were recorded (550 nm; 30°C) (e = 19.2 mM^-1^ ·cm^-1^) after the addition of 20 mM succinate and 10 μM antimycin A. For complex IV (cytochrome *c* oxidase) activity, the first rate constant of cytochrome *c* oxidation was determined^46^ in the presence of 10 mM phosphate buffer and 50 μM reduced cytochrome *c*; absorbance was recorded every minute at 550 nm, 30 °C (e = 19.2 mM^-1^ ·cm^-1^). Citrate synthase activity^47^ was measured in the presence of 93 mM Tris-HCl, 0.1% (vol/vol) Triton X-100, 0.2 mM acetyl-CoA, 0.2 mM DTNB; the reaction was started with 0.2 mM oxaloacetate, and the absorbance was recorded at 412 nm (30 °C) (e = 13.6 mM^-1^ ·cm^-1^).

### Protein Determinations

Protein samples were quantified by the BCA protein assay kit (Thermo) following the manufacturer’s instructions, using BSA as a standard.

### Stereotaxic cannula implantation

For intra-ventricular injections (ICV) a cannula needed to be placed following methods previously published^48,49^. Anesthetized mice with sevoflurane (Sevorane; Abbott) were placed in a stereotaxic frame (Model 1900; David Kopf Instruments) with a micromanipulator (Model 1940; David Kopf Instruments) and a digital reading system (Wizard 550; Anilam). Cranium was exposed to properly reach the coordinates for the lateral ventricle (coordinates from bregma: anteroposterior:-0,2; centre-centre: ±0,9; dorsoventral: −2)^50^, where a small skull hole was perform with a drill (Model 1911; Kopf Instruments). The cannula was placed (C315G/SPC; Plastics-One) and fixed with glue (Loctite 454; Henkel) and dentist cement (Hiflex RR; PrevestDenPro). A dummy was placed (C315GS-5-SP; Plastics-One) to prevent dust brain contamination. Mice after surgery were kept above heating plates (Plactronic Digital; JP Selecta) and feed with soaked food until recovery (at least for 15 days).

### Pharmacokinetics of AZ67

For pharmacokinetic assay healthy male C57BL/6 mice were used. A single dose of 40 mg/kg of AZ67 was injected intravenously and the blood, cerebrospinal fluid (CSF) and brain, were collected after 5 min, 15 min, 30 min, 1 h, 2 h, 4 h, 8 h and 24 h. AZ67 concentration in the different samples were determined by liquid chromatography followed by MS/MS^34^.

### *In vivo* toxicity assay

Male mice (C57BL6/J; 6 animals per group; 8 weeks-old) (purchased from Charles-River, Spain) were subjected to the implantation of a canula in the lateral ventricle under anaesthesia and then left for at least 15 days for fully recovery. After this, the PFKFB3 inhibitor (AZ67) was administered through the canula using an automatic micro-pump (CMA 4004 Microdialysis Syringe Pump, Harvard Apparatus) at different doses: 0 (vehicle), 0.005, 0.01, 0.05, 0.1, 1 and 10 nmol/mice. The compounds were administered every 24 hours for one week and animals were analyzed in the open field immediately before each administration. We selected the maximal dose that caused no evident alterations and/or deterioration of the animals for the following experiments, being 1 nmol/mice.

### Open field tests

Male mice were left to acclimate in the room for no less than 15 min at the same time of day (10:00 to 14:00). Tracking was carried out one at a time, and we carefully cleaned the apparatus with 70% ethanol between trials to remove any odor cues. An ANY-box core was used, which contained a light-grey base and an adjustable perpendicular stick holding a camera and an infrared photo-beam array to track the animal movement and to detect rearing behaviour, respectively. Mouse movements were tracked with the ANY-maze software and the ANY-maze interface to register all parameters described subsequently. For the open field test, a 40 cm × 40 cm × 35 cm (w, d, h), black infrared transparent Perspex insert was used, and the arena was divided in three zones, namely border (8 cm wide), centre (16% of total arena) and intermediate (the remaining area). The test lasted for 10 min, and the distance travelled, and the time spent in each zone were measured.

### AZ67 *in vivo* administration

AZ67 (Tocris) for *in vivo* usage was dissolved in 20% (wt/vol) PEG200 in PBS to a 20 mM concentration. 4 groups were generated (4-6 animals/group), namely: WT-vehicle, CLN7^Δex2^-vehicle, WT-AZ67, CLN7^Δex2^-AZ67. The canula was inserted intracerebroventricularly at the age of 8 weeks and, after at least of 15 days of recovery, we injected the AZ67 at the dose identified previously (1 nmol/mouse) every 24 h. The duration of the experiment was determined by the presence of hindlimb clasping the CLN7^Δex2^-control (vehicle), being this time two months. After this, the animals were perfused, and their brains dissected to be investigated by immunofluorescence and electron microscopy (EM).

### Electron microscopy and mitochondrial morphology analysis

Male mice were anaesthetized by intraperitoneal injection of a mixture of xylazine hydrochloride (Rompun; Bayer) and ketamine hydrochloride/chlorbutol (Imalgene; Merial) (1:4) at 1 ml per kg body weight and then perfused intra-aortically with 0.9% NaCl followed by 5 ml/g body weight of 2% (wt/vol) paraformaldehyde *plus* 2% (vol/vol) glutaraldehyde. After perfusion, brains were dissected out sagitally in two parts and post-fixed with perfusion solution over night at 4°C. Brain blocks were rinsed with 0.1 M PB solution and a 1 mm^3^ squared of brain cortex was excised and treated with osmium tetroxide (1% in PB) for 1 h. Tissue was then washed with distilled water, and dehydrated in ascending series of ethanol followed by embedment in EPON resin. Ultra-thin sections (50 nm) were stained with uranyl acetate and lead citrate and examined with Tecnai Spirit Twin 120 kv transmission electron microscopy equipped with a digital camera Orius WD or JEM-1010 (JEOL) 100 kv transmission electron microscopy equipped with a digital camera AMT RX80. For mitochondrial area quantification, the area of each mitochondria was quantified in neuronal soma, axons and dendrites. In the case of mitochondrial length, the values represent the length in the maximal axis of mitochondria in the plane of microphotographies. Cristae profiles of representative mitochondria of each condition and type were traced along the major axis that crosses mitochondria perpendicularly to cristae. Data of pixel intensity were obtained using the plot profile plugin of ImageJ software.

### Mouse perfusion and immunohistochemistry

Mice (5 months for AZ67 intraventricular injections; 3 months for mCAT expression approach) were anaesthetized by intraperitoneal injection of a mixture of xylazine hydrochloride (Rompun; Bayer) and ketamine hydrochloride/chlorbutol (Imalgene; Merial) (1:4) at 1 ml per kg body weight and then perfused intra-aortically with 0.9% NaCl followed by 5 ml p/g body weight of Somogyi (4% (wt/vol) paraformaldehyde, and 0.2% (vol/vol) picric acid, in 0.1 M PB; pH 7.4). After perfusion, brains were dissected out sagitally in two parts and post-fixed with Somogyi for 2 h at room temperature. Brain blocks were rinsed successively for 10 min, 30 min and 2 h with 0.1 M PB solution and cryoprotected in 10%, 20% and 30% (wt/vol) sucrose in PB sequentially, until they sank. After cryoprotection, 40-μm-thick sagittal sections were obtained with a freezing-sliding cryostat (Leica). Sectioning of WT and *Cln7*^Δ*ex2*^ brains were performed under the same conditions and sessions. The sections were collected serially in a 12-well plate in 0.1 M PB, rinsed three times for 10 min in 0.1 M PB and used for subsequent immunohistochemistry and lipofuscin observation. The section-containing wells that were not used were kept in freezer mix (30% (vol/vol) polyethylene glycol, 30% (vol/vol) glycerol in 0.1 M PB) at −20°C. For immunohistochemistry, sections were incubated sequentially in (i) 5 mg/ml sodium borohydride in PB for 30 min (to remove aldehyde autofluorescence); (ii) three PBS washes of 10 min each; (iii) 1/500 anti-GFAP (G6171; Sigma) and 1/500 anti-IBA1 (019–19741; Wako) or 1/500 anti-ATP-C (SCMAS) (ab181243; Abcam) in 0.02% Triton X-100 (Sigma) and 5% goat serum (Jackson Immuno-Research) in 0.1 M PB for 72 h at 4 °C; (iv) three PB washes of 10 min each; (v) fluorophore conjugated secondary antibodies, 1/500 Cy2 goat anti-mouse and 1/500 Cy3 goat anti-rabbit (Jackson Immuno-Research) or Alexa-488 (A11008; Molecular Probes) in PB for 2 h at room temperature; and (vi) 0.5 μg/ml DAPI in PB for 10 min at room temperature. After being rinsed with PB, sections were mounted with Fluoromount (Sigma) aqueous mounting medium and cover slips (Thermo Fisher)^51^. For autofluorescence (lipofuscine accumulation) sections were mounted directly.

### Imaging and quantification

Sections were examined with epifluorescence and the appropriate filter sets under an Operetta CLS high-content imaging system (PerkinElmer). Large fields of view were acquired with an 5x scan using an OperaPHX/OPRTCLS 5x Air Objective. Then high-resolution images were acquired using an OperaPHX/OPRTCLS Air Objective 20x hNA objective. Immunohistochemical digital images were used to analyse different proteins staining in the three most sagittal sections per animal from, at least, two different animals per condition (n=2). Images were analysed with the Harmony software with PhenoLOGIC (PerkinElmer). Interest brain areas (cortex, hippocampus and cerebellum) were selected and subsequently quantified as mean intensity per area by using the ‘measure rectangle’ function, which represents the mean intensity of a channel per selected area.

### NPC immunocytochemistry

NPCs were fixed with 100% iced-cold methanol for 5 min and incubated in blocking solution (1% (v/v) normal goat serum, 0.1% (w/v) bovine serum albumin (BSA), 0.1% (v/v) Triton X-100 in DPBS). The antibodies were incubated in blocking solution. The incubation of the primary antibody (mouse α-ATP5A, 1:100, Abcam, ab14748) or SCMAS (1/200, ab181243, Abcam) was performed for 2 h at room temperature, and the secondary antibodies (Alexa Fluor 568 goat α-mouse, 1:500, Invitrogen or Alexa Fluor 488 goat anti-rabbit, A-11008, Thermo) were applied for 1 h at room temperature. Slides were mounted with VECTASHIELD Mounting Medium with DAPI, incubated for 24 h at 4 °C and imaged with a Zeiss Axio Imager M2 fluorescence microscope or under an inverted microscope (Nikon; Eclipse Ti-E) equipped with a pre-centred fibre illuminator (Nikon; Intensilight C-HGFI), B/W CCD digital camera (Hamamatsu; ORCA-E.R.). Fluorescence quantification was performed after appropriate thresholding using the ImageJ software (NIH). The pixel intensity profile of ATP5A immunodecoration was analyzed across the maximal axis of the cell that departs from the nucleus, using the plot profile plugin of ImageJ software. A representative profile is shown for each condition.

### Statistical Analysis

The comparisons between two groups of values we performed Student’s *t* test. For multiple-values comparisons we used one-way ANOVA, post-hoc Tukey or DMS. The statistical analysis was performed using the GraphPad Prism v8 software. The number of biologically independent culture preparations or animals used per experiment and the *P* values are indicated in the figure legends.

